# Environmental and epigenetic regulation of *Rider* retrotransposons in tomato

**DOI:** 10.1101/517508

**Authors:** Matthias Benoit, Hajk-Georg Drost, Marco Catoni, Quentin Gouil, Sara Lopez-Gomollon, David Baulcombe, Jerzy Paszkowski

## Abstract

Transposable elements in crop plants are the powerful drivers of phenotypic variation that has been selected during domestication and breeding programs. In tomato, transpositions of the LTR (long terminal repeat) retrotransposon family *Rider* have contributed to various phenotypes of agronomical interest, such as fruit shape and colour. However, the mechanisms regulating *Rider* activity are largely unknown. We have developed a bioinformatics pipeline for the functional annotation of retrotransposons containing LTRs and defined all full-length *Rider* elements in the tomato genome. Subsequently, we showed that accumulation of *Rider* transcripts and transposition intermediates in the form of extrachromosomal DNA is triggered by drought stress and relies on abscisic acid signalling. We provide evidence that residual activity of *Rider* is controlled by epigenetic mechanisms involving siRNAs and the RNA-dependent DNA methylation pathway. Finally, we demonstrate the broad distribution of *Rider-like* elements in other plant species, including crops. Thus our work identifies *Rider* as an environment-responsive element and a potential source of genetic and epigenetic variation in plants.

## INTRODUCTION

Transposable elements (TEs) replicate and move within host genomes. Based on their mechanisms of transposition, TEs are either DNA transposons that use a cut-and-paste mechanism or retrotransposons that transpose through an RNA intermediate via a copy-and-paste mechanism [1]. TEs make up a significant part of eukaryotic chromosomes and are a major source of genetic instability that, when active, can induce deleterious mutations. Various mechanisms have evolved that protect plant genomes, including the suppression of TE transcription by epigenetic silencing that restricts TE movement and accumulation [2–5].

Chromosomal copies of transcriptionally silenced TEs are typically hypermethylated at cytosine residues and are associated with nucleosomes containing histone H3 di-methylated at lysine 9 (H3K9me2). In addition, they are targeted by 24-nt small interfering RNAs (24-nt siRNAs) that guide RNA-dependent DNA methylation (RdDM), forming a self-reinforcing silencing loop [6–8]. Interference with these mechanisms can result in the activation of transposons. For example, loss of DNA METHYLTRANSFERASE 1 (MET1), the main methyltransferase maintaining methylation of cytosines preceding guanines (CGs), results in the activation of various TE families in Arabidopsis [9–11] and in rice [12]. Mutation of CHROMOMETHYLASE 3 (CMT3), mediating DNA methylation outside CGs, triggers the mobilization of several TE families, including *CACTA* elements in *Arabidopsis* [10] and *Tos17* and *Tos19* in rice [13]. Interference with the activity of the chromatin remodelling factor DECREASE IN DNA METHYLATION 1 (DDM1), as well as various components of the RdDM pathway, leads to the activation of specific subsets of TEs in Arabidopsis. These include DNA elements *CACTA* and *MULE*, as well as retrotransposons *ATGP3*, *COPIA13*, *COPIA21*, *VANDAL21*, *EVADÉ* and *DODGER* [14–17]. Similarly, loss of *OsDDM1* genes in rice results in the transcriptional activation of TE-derived sequences [18].

In addition to interference with epigenetic silencing, TE activation can also be triggered by environmental stresses. In her pioneering studies, Barbara McClintock denoted TEs as “controlling elements”, thus suggesting that they are activated by genomic stresses and are able to regulate the activities of genes [19, 20]. In the meantime, a plethora of stress-induced TEs have been described, including retrotransposons. For example, the biotic stress-responsive *Tnt1* and *Tto1* families in tobacco [21,22], the cold-responsive *Tcs* family in citrus [23], the virus-induced *Bs1* retrotransposon in maize [24], the heat-responsive retrotransposons *Go-on* in rice [25], and *ONSEN* in Arabidopsis [26,27]. While heat-stress is sufficient to trigger *ONSEN* transcription and the formation of extrachromosomal DNA (ecDNA), transposition was observed only after the loss of siRNAs, suggesting that the combination of impaired epigenetic control and environmental stress is a prerequisite for *ONSEN* transposition [28]. Interestingly, retrotransposition occurs during flower development, which fuels the diversification of *ONSEN* insertion patterns in the progenies of plants permitting *ONSEN* movement [29].

The availability of high-quality genomic sequences revealed that LTR (Long Terminal Repeat) retrotransposons make up a significant proportion of plant chromosomes, from approximately 10% in Arabidopsis, 25% in rice, 42% in soybean, and up to 75% in maize [30]. In tomato (*Solanum lycopersicum*), a model crop plant for research on fruit development, LTR retrotransposons make up about 60% of the genome [31]. Despite the abundance of retrotransposons in the tomato genome, only a limited number of studies have linked TE activities causally to phenotypic alterations. Remarkably, the most striking examples described so far involve the retrotransposon family *Rider*. For example, fruit shape variation is based on copy number variation of the *SUN* gene, which underwent *Rider*-mediated trans-duplication from chromosome 10 to chromosome 7. The new insertion of the *SUN* gene into chromosome 7 in the variety “Sun1642” results in its overexpression and consequently in the elongated tomato fruits that were subsequently selected by breeders [32,33]. The *Rider* element generated an additional *SUN* locus on chromosome 7 that encompassed more than 20 kb of the ancestral *SUN* locus present on chromosome 10 [32]. This large “hybrid” retroelement landed in the fruit-expressed gene *DEFL1*, resulting in high and fruit-specific expression of the *SUN* gene containing the retroelement [33]. The transposition event was estimated to have occurred within the last 200-500 years, suggesting that duplication of the *SUN* gene occurred after tomato domestication [34].

Jointless pedicel is a further example of a *Rider*-induced tomato phenotype that has been selected during tomato breeding. This phenotypic alteration reduces fruit dropping and thus facilitates mechanical harvesting. Several independent jointless alleles were identified around 1960 [35–37]. One of them involves a new insertion of *Rider* into the first intron of the *SEPALLATA* MADS-Box gene, *Solyc12g038510*, that provides an alternative transcription start site and results in an early nonsense mutation [38]. Also, the ancestral yellow flesh mutation in tomato is due to *Rider*-mediated disruption of the *PSY1* gene, which encodes a fruit-specific phytoene synthase involved in carotenoid biosynthesis [39,40]. Similarly, the “potato leaf” mutation is due to a *Rider* insertion in the *C* locus controlling leaf complexity [41]. *Rider* retrotransposition is also the cause of the chlorotic tomato mutant *fer*, identified in the 1960s [42]. This phenotype has been linked to *Rider*-mediated disruption of the *FER* gene encoding a bHLH-transcription factor. *Rider* landed in the first exon of the gene [43,44]. Sequence analysis of the element revealed that the causative copy of *Rider* is identical to that involved in the *SUN* gene duplication [44].

The *Rider* family belongs to the *Copia* superfamily and is ubiquitous in the tomato genome [33,44]. Based on partial tomato genome sequences, the number of *Rider* copies was estimated to be approximately 2000 [33]. Previous DNA blots indicated that *Rider* is also present in wild tomato relatives but is absent from the genomes of potato, tobacco, and coffee, suggesting that amplification of *Rider* happened after the divergence of potato and tomato approximately 6.2 mya [44,45]. The presence of *Rider* in unrelated plant species has also been suggested [46]. However, incomplete sub-optimal sampling and the low quality of genomic sequence assemblies has hindered a comprehensive survey of *Rider* elements within the plant kingdom.

Considering that the *Rider* family is a major source of phenotypic variation in tomato, it is surprising that its members and their basic activities, as well as their responsiveness and the possible triggers of environmental super-activation, which explain the evolutionary success of this family, remain largely unknown. Contrary to the majority of TEs characterized to date, previous analyses revealed that *Rider* is constitutively transcribed and produces full-length transcripts in tomato [33], but the stimulatory conditions promoting reverse transcription of *Rider* transcripts that results in accumulation as extrachromosomal DNA are unknown.

To fill these gaps, we provide here a refined annotation of full-length *Rider* elements in tomato using the most recent genome release (SL3.0). We reveal environmental conditions facilitating *Rider* activation and show that *Rider* transcription is enhanced by dehydration stress mediated by abscisic acid (ABA) signalling, which also triggers accumulation of extrachromosomal DNA. Moreover, we provide evidence that RdDM controls *Rider* activity through siRNA production and partially through DNA methylation. Finally, we have performed a comprehensive cross-species comparison of full-length *Rider* elements in 110 plant genomes, including diverse tomato relatives and major crop plants, in order to characterise species-specific *Rider* features in the plant kingdom. Together, our findings suggest that *Rider* is a drought stress-induced retrotransposon ubiquitous in diverse plant species that may have contributed to phenotypic variation through the generation of genetic and epigenetic alterations induced by historical drought periods.

## MATERIAL AND METHODS

### Plant material and growth conditions

Tomato plants were grown under standard greenhouse conditions (16 h at 25°C with supplemental lighting of 88 w/m^2^ and 8 h at 15°C without). *flacca* (*flc*), *notabilis* (*not*), and *sitiens* (*sit*) seeds were obtained from Andrew Thompson, Cranfield University; the *slnrpd1* and *slnrpe1* plants were described before [47]. For aseptic growth, seeds of *Solanum lycopersicum* cv. Ailsa Craig were surface-sterilized in 20% bleach for 10 min, rinsed three times with sterile H_2_O, germinated and grown on half-strength MS media (16 h light and 8 h dark at 24°C).

### Stress treatments

For dehydration stress, two-week-old greenhouse-grown plants were subjected to water deprivation for two weeks. For NaCl and mannitol treatments, tomato seedlings were grown aseptically for two weeks prior to transfer into half-strength MS solution containing 100, 200 or 300 nM NaCl or mannitol (Sigma) for 24 h. For abscisic acid (ABA) treatments, tomato seedlings were grown aseptically for two weeks prior to transfer into half-strength MS solution containing 0.5, 5, 10 or 100 μM ABA (Sigma) for 24 h. For 5-azacytidine treatments, tomato seedlings were germinated and grown aseptically on half-strength MS media containing 50 nM 5-azacytidine (Sigma) for two weeks. For cold stress experiments, two-week-old aseptically grown plants were transferred to 4°C for 24 h prior to sampling.

### RNA extraction and quantitative RT-PCR analysis

Total RNA was extracted from 200 mg quick-frozen tissue using the TRI Reagent (Sigma) according to the manufacturer’s instructions and resuspended in 50 μL H_2_O. The RNA concentration was estimated using the Qubit Fluorometric Quantitation system (Thermo Fisher). cDNAs were synthesized using a SuperScript VILO cDNA Synthesis Kit (Invitrogen). Real-time quantitative PCR was performed in the LightCycler 480 system (Roche) using primers listed in Table S1. LightCycler 480 SYBR Green I Master premix (Roche) was used to prepare the reaction mixture in a volume of 10 μL. Transcript levels were normalized to *SlACTIN* (*Solyc03g078400*). The results were analysed by the ΔΔCt method.

### DNA extraction and copy number quantification

Tomato DNA was extracted using the Qiagen DNeasy Plant Mini Kit (Qiagen) following the manufacturer’s instructions and resuspended in 30 μL H_2_O. DNA concentration was estimated using the Qubit Fluorometric Quantitation system (Thermo Fisher). Quantitative PCR was performed in the LightCycler 480 system (Roche) using primers listed in Table S1. LightCycler 480 SYBR Green I Master premix (Roche) was used to prepare the reaction in a volume of 10 μL. DNA copy number was normalized to *SlACTIN* (*Solyc03g078400*). Results were analysed by the ΔΔCt method.

### Extrachromosomal circular DNA detection

Extrachromosomal circular DNA amplification was derived from the previously published mobilome analysis [11]. In brief, extrachromosomal circular DNA was separated from chromosomal DNA using PlasmidSafe ATP-dependent DNase (EpiCentre) according to the manufacturer’s instructions with the incubation at 37°C extended to 17 h. The PlasmidSafe exonuclease degrades linear DNA and thus safeguards circular DNA molecules. Circular DNA was precipitated overnight at −20°C in 0.1 v/v 3 M sodium acetate (pH 5.2), 2.5 v/v EtOH and 1 μL glycogen (Sigma). The pellet was resuspended in 20 μL H_2_O. Inverse PCR reactions were carried out with 2 μL of DNA solution in a final volume of 20 μL using the GoTaq enzyme (Promega). The PCR conditions were as follows: denaturation at 95°C for 5 min, followed by 30 cycles at 95°C for 30 s, an annealing step for 30 s, an elongation step at 72°C for 60 s, and a final extension step at 72°C for 5 min. PCR products were separated in 1% agarose gels and developed by NuGenius (Syngene). Bands were extracted using the Qiagen Gel Extraction Kit and eluted in 30 μL H_2_O. Purified amplicons were subjected to Sanger sequencing. Primer sequences are listed in Table S1.

### Phylogenetic analysis of de novo identified Rider elements

A phylogenetic tree was constructed from the nucleotide sequences of the 71 *Rider* elements using Geneious 9.1.8 (www.geneious.com) and built with the Tamura-Nei neighbor joining method. Pairwise alignment for the building distance matrix was obtained using a global alignment with free end gaps and a cost matrix of 51% similarity.

### Distribution analysis

Genomic coordinates of each of the 71 *Rider* elements identified by *de novo* annotation using *LTRpred* (https://github.com/HajkD/LTRpred) have been used to establish their chromosomal locations. Coordinates for centromeres were provided before [31] and pericentromeric regions were defined by high levels of DNA methylation and H3K9me2 ([47] and David Baulcombe, personal communication).

### Accession numbers

The Genbank accession number of the reference *Rider* nucleotide sequence identified in [44] is EU195798.2. We used *Solanum lycopersicum* bisulfite and small RNA sequencing data (SRP081115) generated in [47].

### Dating of insertion time

Insertion times of *Rider* elements were estimated using the method described in [44]. Degrees of divergence between LTRs of each individual element were determined using *LTRpred*. LTR divergence rates were then converted into dates using the average substitution rate of 6.96 x 10^-9^ substitutions per synonymous site per year for tomato [48].

### Bisulfite sequencing analysis

We collected data from previously published BS-seq libraries of tomato mutants of RNA polymerase IV and V and controls [47]: *slnrpe1* (SRR4013319), *slnrpd1* (SRR4013316), wild type *CAS9* (SRR4013314) and not transformed wild type (SRR4013312). The raw reads were analysed using our previously established pipeline [49] and aligned to the *Solanum lycopersicum* reference version SL3.0 (www.solgenomics.net/organism/Solanum_lycopersicum/genome). The chloroplast sequence (NC_007898) was used to estimate the bisulfite conversion (on average above 99%). The R package DMRcaller [50] was used to summarize the level of DNA methylation in the three cytosine contexts for each *Rider* copy.

### Small RNA sequencing analysis

Tomato siRNA libraries were obtained from [47] and analysed using the same analysis pipeline to align reads to the tomato genome version SL3.0. Briefly, the reads were trimmed with Trim Galore! (www.bioinformatics.babraham.ac.uk/projects/trim_galore) and mapped using the ShortStack software v3.6 [51]. The siRNA counts on the loci overlapping *Rider* copies were calculated with R and the package GenomicRanges.

### Genome sequence data

Computationally reproducible analysis and annotation scripts for the following sections can be found at http://github.com/HajkD/RIDER.

### Genomic data retrieval

We retrieved genome assemblies for 110 plant species (Table S2) from NCBI RefSeq [52] using the *meta.retrieval* function from the R package *biomartr* [53]. For *Solanum lycopersicum*, we retrieved the most recent genome assembly version SL3.0 from the *Sol Genomics Network* ftp://ftp.solgenomics.net/tomato_genome/assembly/build_3.00/S_lycopersicu *m_chromosomes.3.00.fa* [54].

### Functional *de novo* annotation of LTR retrotransposons in *Solanaceae* genomes

Functional *de novo* annotations of LTR retrotransposons for seventeen genomes from the *Asterids, Rosids*, and *monocot* clades (*<u>Asterids</u>*: *Capsicum annuum*, *C. baccatum MLFT02_5*, *C. chinense MCIT02_5*, *Coffea canephora*, *Petunia axillaris*, *Phytophthora inflata*, *Solanum arcanum*, *S. habrochaites*, *S. lycopersicum*, *S. melongena*, *S. pennellii*, *S. pimpinellifolium*, *S. tuberosum*; *<u>Rosids</u>*: *Arabidopsis thaliana*, *Vitis vinifera*, and *Cucumis melo*; *<u>Monocots</u>*: *Oryza sativa*) were generated using the *LTRpred.meta* function from the *LTRpred* annotation pipeline (https://github.com/HajkD/LTRpred; also used in [25]). To retrieve a consistent and comparable set of functional annotations for all genomes, we consistently applied the following *LTRpred* parameter configurations to all *Solanaceae* genomes: minlenltr = 100, maxlenltr = 5000, mindistltr = 4000, maxdisltr = 30000, mintsd = 3, maxtsd = 20, vic = 80, overlaps = “no”, xdrop = 7, motifmis = 1, pbsradius = 60, pbsalilen = c(8,40), pbsoffset = c(0,10), quality.filter = TRUE, n.orf = 0. The plant-specific tRNAs used to screen for primer binding sites (PBS) were retrieved from GtRNAdb [55] and plant RNA [56] and combined in a custom *fasta* file. The hidden Markov model files for gag and pol protein conservation screening were retrieved from Pfam [57] using the protein domains RdRP_1 (PF00680), RdRP_2 (PF00978), RdRP_3 (PF00998), RdRP_4 (PF02123), RVT_1 (PF00078), RVT_2 (PF07727), Integrase DNA binding domain (PF00552), Integrase zinc binding domain (PF02022), Retrotrans_gag (PF03732), RNase H (PF00075), and Integrase core domain (PF00665).

### Sequence clustering of functional LTR retrotransposons from 17 genomes

We combined the *de novo* annotated LTR retrotransposons of the 17 species mentioned in the previous section in a large fasta file and used the cluster program *VSEARCH* [58] with parameter configurations: *vsearch --cluster_fast --qmask none –id 0.85 --clusterout_sort --clusterout_id --strand both -- blast6out ---sizeout* to cluster LTR retrotransposons by nucleotide sequence homology (global sequence alignments). Next, we retrieved the 85% sequence homology clusters from the *VSEARCH* output and screened for clusters containing *Rider* sequences. This procedure enabled us to detect high sequence homology (>85%) sequences of *Rider* across diverse species.

### Nucleotide BLAST search of *Rider* against 110 plant genomes

To determine the distribution of *Rider* related sequences across the plant kingdom, we performed BLASTN [59] searches of *Rider* (= query sequence) using the function *blast_genomes* from the R package *metablastr* (https://github.com/HajkD/metablastr) against 110 plant genomes (Table S2) and the parameter configuration: *blastn -eval 1E-5 -max_target_seqs 5000*. As a result, we retrieved a BLAST hit table containing 11,748,202 BLAST hits. Next, we filtered for hits that contained at least 50% sequence coverage (= sequence homology) and throughout at least 50% sequence length homology to the reference *Rider* sequence. This procedure reduced the initial 11,748,202 BLAST hits to 57,845 hits, which we further refer to as *Rider-like* elements. These 57,845 *Rider-like* elements are distributed across 21 species with various abundance frequencies. In a second step, we performed an analogous BLASTN search using only the 5’ LTR sequence of *Rider* to determine the distribution of *Rider-like* LTR across the plant kingdom. Using the same BLASTN search strategy described above, we retrieved 9,431 hits. After filtering for hits that contained at least 50% percent sequence coverage (= sequence homology) and at least 50% sequence length homology to the reference *Rider* LTR sequence, we obtained 2,342 BLAST hits distributed across five species.

### Motif enrichment analysis

We tested the enrichment of *cis*-regulatory elements (CREs) in *Rider* using two approaches. In the first approach, we compared *Rider* CREs to promoter sequences of all 35,092 protein coding genes from the tomato reference genome. We retrieved promoter sequences 400 bp upstream of the TSS of the respective genes. We constructed a 2×2 contingency table containing the respective motif count data of CRE observations in true *Rider* sequences *versus* counts in promoter sequences. We performed a Fisher’s exact test for count data to assess the statistical significance of enrichment between the motif count data retrieved from *Rider* sequences and the motif count data retrieved from promoter sequences. In the second approach, due to the unavailability of gene annotation for *Solanum arcanum*, *Solanum habrochaites* and *Solanum pimpinellifolium* we compared *Rider* CREs to randomly sampled sequence loci from the same genome using the following two step procedure: in step one, we sampled 1000 DNA sequences with the same length as the reference *Rider* sequence from 1000 randomly sampled loci in the tomato reference genome. When sampling, we also considered the strand direction of the reference *Rider* sequence. Whenever a *Rider* sequence was annotated in the plus direction, we also sampled the corresponding set of random sequences in the plus direction of the respective randomly drawn locus. In contrast, when a *Rider* sequence was annotated in the minus direction, we also sampled the corresponding set of random sequences in the minus direction. In step two, we counted CRE occurrences for each *Rider* sequence independently and for a set of different CREs. Next, we counted the number of the same CRE occurrences for each random sequence independently to assess how often these CREs were found in random sequences. We then, analogous to the first approach, constructed a 2×2 contingency table containing the respective motif count data of CRE observations in true *Rider* sequences *versus* counts in random sequences. We performed a Fisher’s exact test for count data to assess the statistical significance of enrichment between the motif count data retrieved from *Rider* sequences and the motif count data retrieved from random sequences. The resulting *P*-values are shown in Table S3 for the first approach and in Table S4 for the second approach. Computationally reproducible scripts to perform the motif count analysis can be found at https://github.com/HajkD/RIDER.

### Calculation of N50 metric

To assess the genome quality of *Solanaceae* species, we calculated the N50 metric for the genome assemblies of *Solanum lycopersicum, S. pimpinellifolium, S. arcanum, S. pennellii, S. habrochaites*, and *S. tuberosum* using the following procedure. First, we imported the scaffolds or chromosomes of each respective genome assembly using the R function *read_genome()* from the *biomartr* package. Next, for each species individually we determined the sequence length for each scaffold or chromosome and sorted them according to length in descending order. The N50 value in Mbp was then calculated in R as follows: *N50 <- len.sorted*[*cumsum(len.sorted) >= sum(len.sorted)*0.5][1]/1000000*, where the variable *len.sorted* denotes the vector storing the ordered scaffold or chromosome lengths of a genome assembly.

## RESULTS

### Family structure of *Rider* retrotransposons in tomato

We used the most recent SL3.0 tomato genome release for *de novo* annotation of *Rider* elements. First, we retrieved full-length, potentially autonomous retrotransposons using our functional annotation pipeline (*LTRpred*, see Materials and Methods). We detected a set of 5844 potentially intact LTR retrotransposons (Table S5). Homology search among these elements identified 71 elements that share >85% similarity with the reference *Rider* sequence [44] and thus belong to the *Rider* family. We then determined the distribution of these *Rider* elements along the tomato chromosomes (Figure 1A) and also estimated their age based on sequence divergence between 5’ and 3’ LTRs (Figure 1A). We classified these elements into three categories according to their LTR similarity: 80-95%, 95-98% and 98-100% (Figure S1A). While the first category contains relatively old copies (last transposition between 10.5 and 3.5 mya), the 95-98% class represents *Rider* elements that moved between 3.5 and 1.4 mya, and the 98-100% category includes the youngest *Rider* copies that transposed within the last 1.4 my (Figure S1A). Out of 71 *Rider* family members, 14 were found in euchromatic chromosome arms (14/71 or 19.7%) and 57 in heterochromatic regions (80.3%) (Table 1). In accordance with previous observations based on partial genomic sequences [33], young *Rider* elements of the 98-100% class are more likely to reside in the proximity of genes, with 50% within 2 kb of a gene. This was the case for only 37.5% of old *Rider* members (85-95% class) (Table 2). Such a distribution is consistent with the preferential presence of young elements within euchromatic chromosome arms (50%, 5/10) compared to old *Rider* elements (9.4%, 3/32) (Table 2 and Figure S1B). In addition, the phylogenetic distance between individual elements is moderately correlated to the age of each element (Figure 1B) (Table S6).

**Figure 1:**
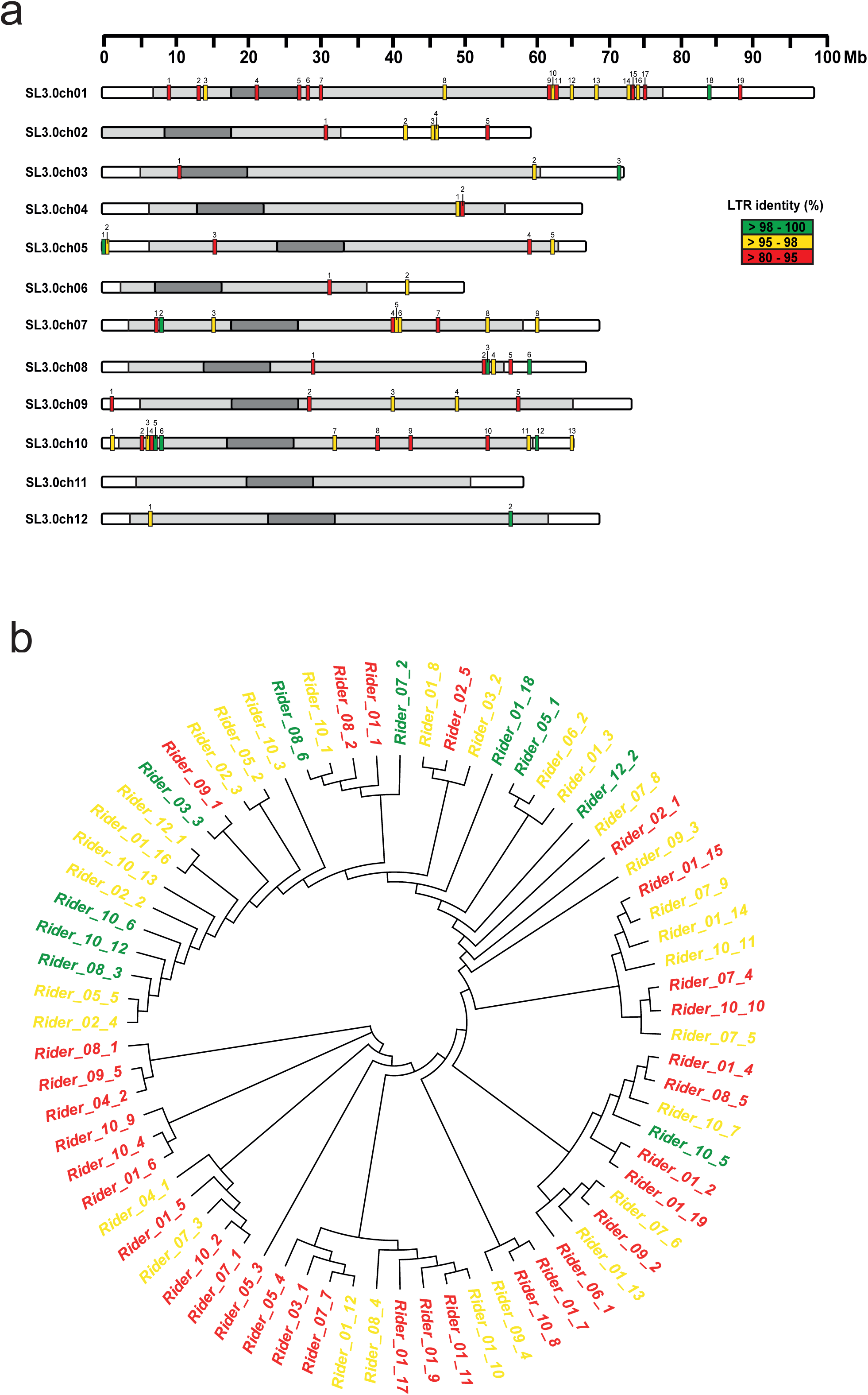
Chromosomal location and phylogenetic relationships of *de novo* annotated full-length *Rider* elements. (A) Chromosomal positions of 71 *de novo* annotated full-length *Rider* elements in the SL3.0 genome. *Rider* copies are marked as coloured vertical bars, with colours reflecting similarity between LTRs for each element. Dark grey areas delimitate the centromeres, light grey pericentromeric heterochromatin, and white euchromatin. (B) Phylogenetic relationship of the 71 *de novo* annotated *Rider* elements. The phylogenetic tree was constructed using the neighbour-joining method on nucleotide sequences of each *Rider* copy.

**Table 1:**
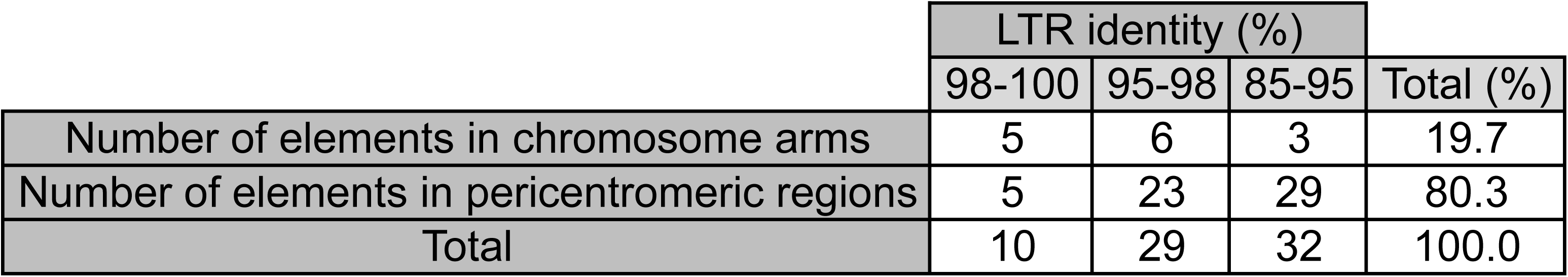
Distribution of *de novo* annotated *Rider* elements based on chromatin context.

**Table 2:** Distribution of *de novo* annotated *Rider* elements based on gene proximity.

|  | Presence of gene within 2 kb (%) | Number of elements in chromosome arms (%) |
| --- | --- | --- |
| Rider_85-95 | 37.5 | 9.4 |
| Rider_95-98 | 48.3 | 20.7 |
| Rider_98-100 | 50.0 | 50 |

### *Rider* is a drought- and ABA-responsive retrotransposon

To better understand the activation triggers and, thus, the mechanisms involved in the accumulation of *Rider* elements in the tomato genome, we examined possible environmental stresses and host regulatory mechanisms influencing their activity. Transcription of an LTR retroelement initiates in its 5’ LTR and is regulated by an adjacent promoter region that usually contains *cis*- regulatory elements (CREs) (reviewed in [60]). Therefore, we aligned the sequence of the *Rider* promoter region against sequences stored in the PLACE database (www.dna.affrc.go.jp/PLACE/) containing known CREs and identified several dehydration-responsive elements (DREs) and sequence motifs linked to ABA signalling (Figure 2A). First, we tested whether these CREs were present and enriched in the LTR promoter sequences of the 71 *de novo* annotated *Rider* elements (Table S7). Comparison of *Rider* LTRs to a set of gene promoter sequences of the same length revealed significant enrichment of several CREs in *Rider* LTRs (Fisher’s exact test *P*<0.001) (Table S3). It is known, for example, that the CGCG sequence motif at position 89-94 (Figure 2A) is recognized by transcriptional regulators binding calmodulin. These are products of signal-responsive genes activated by various environmental stresses and phytohormones such as ABA [61]. We also detected two MYB recognition sequence motifs (CTGTTG at position 176-181 bp, and CTGTTA at position 204-209 bp) (Figure 2A). MYB recognition sequences are usually enriched in the promoters of genes with transcriptional activation during water stress, elevated salinity, and ABA treatments [62,63]. In addition, an ABA-responsive element-like (ABRE-like) was found at position 332-337 bp in the R region of *Rider*’s LTR, along with a coupling element (CE3) located at position 357-372 bp (Figure 2A). The co-occurrence of ABRE-like and CE3 has often been found in ABA-responsive genes [64,65].

**Figure 2:**
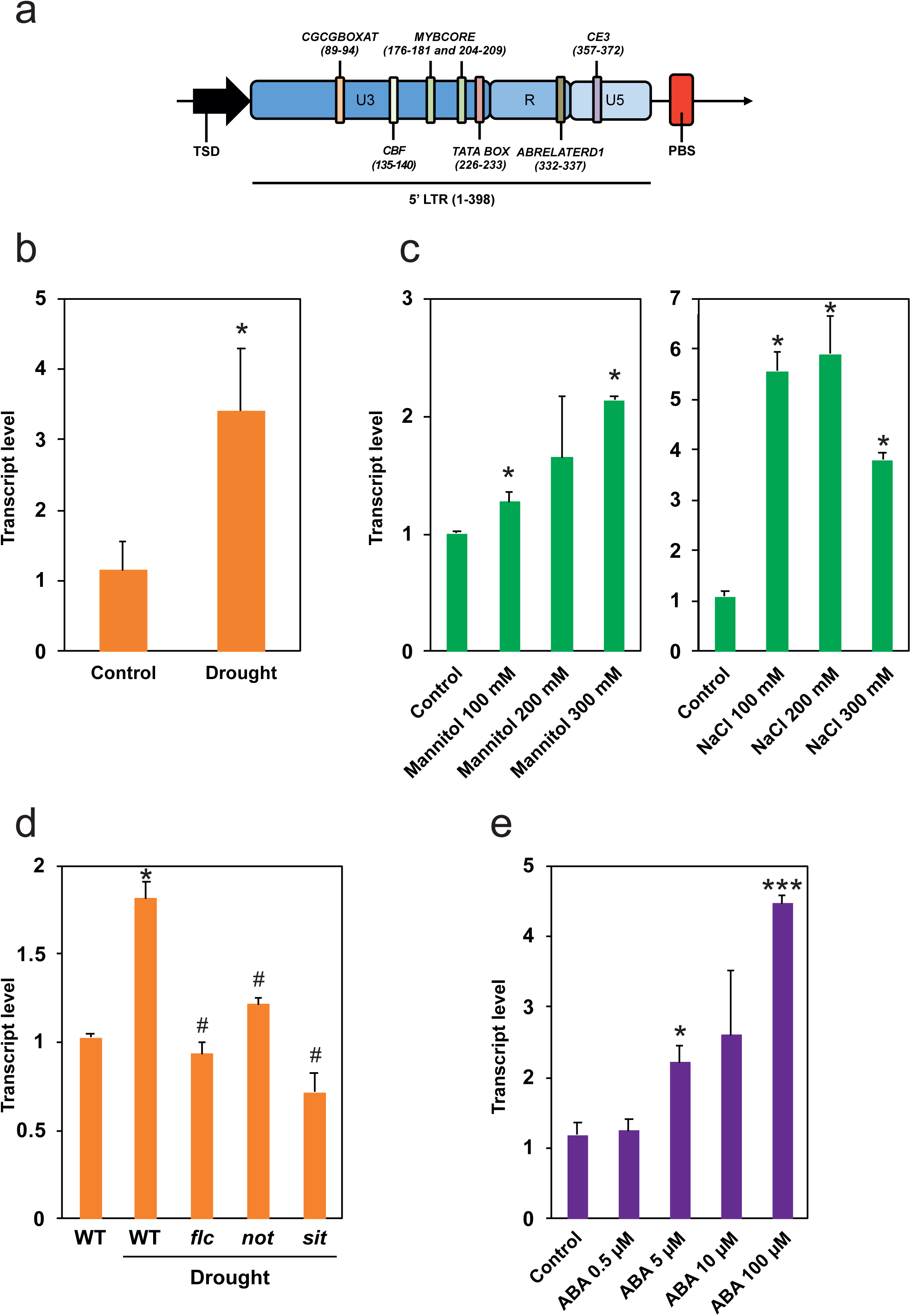
*Rider* activation is stimulated by drought and ABA. (A) Identification of *cis*-regulatory elements (CREs) within *Rider* LTRs. *Rider* LTR U3, R and U5 regions are marked, as well as neighbouring Target Site Duplication (TSD) and Primer Binding Site (PBS) sequences. CREs are marked as coloured vertical bars; their bp positions are given in brackets. (B-C) Quantification of *Rider* RNA levels by RT-qPCR in tomato seedlings after (B) drought stress or (C) mannitol and NaCl treatments. Histograms show normalized expression relative to Control, +/- SEM from two to three biological replicates. \**P*<0.05, two-sided Student’s *t*-test. (D) Quantification of *Rider* RNA levels by RT-qPCR in leaves of drought-stressed tomato wild-type plants, *flc*, *not* and *sit* mutants. Histograms show normalized expression relative to WT Control, +/- SEM from two biological replicates. \**P*<0.05 denotes difference compared to wild-type control; ^#^ *P*<0.05 denotes difference compared to wild-type drought plants, two-sided Student’s *t*-test. (E) Quantification of *Rider* RNA levels by RT-qPCR in tomato seedlings after ABA treatment. Histograms show normalized expression relative to Control, +/- SEM from two to three biological replicates. \**P*<0.05, \*\*\**P*<0.001, two-sided Student’s *t*-test.

The simultaneous presence of these five CREs in promoters of *Rider* elements suggests that *Rider* transcription may be induced by environmental stresses such as dehydration and salinity that involves ABA mediated signalling. To test whether *Rider* transcription is stimulated by drought stress, glasshouse-grown tomato plants were subjected to water deprivation and levels of *Rider* transcripts quantified by RT-qPCR (Figure 2B). When compared to control plants, we observed a 4.4-fold increase in *Rider* transcript abundance in plants subjected to drought stress. Thus, *Rider* transcription appears to be stimulated by drought.

To further test this finding, we re-measured levels of *Rider* transcripts in different experimental setups. *In vitro* culture conditions with increasing levels of osmotic stress were used to mimic increasing drought severity (Figure 2C). Transcript levels of *Rider* increased in a dose-dependent fashion with increasing mannitol concentration, corroborating results obtained during direct drought stress in greenhouse conditions. Interestingly, tomato seedlings treated with NaCl also exhibited increased levels of *Rider* transcripts (Figure 2C).

ABA is a versatile phytohormone involved in plant development and abiotic stress responses, including drought stress [66]. Therefore, we asked whether *Rider* transcriptional drought-responsiveness is mediated by ABA and whether increased ABA can directly stimulate *Rider* transcript accumulation. To answer the first question, we exploited tomato mutants defective in ABA biosynthesis. The lines *flacca* (*flc*), *notabilis* (*not*) and *sitiens* (*sit*) have mutations in genes encoding a sulphurylase [67], a 9-cis-epoxy-carotenoid dioxygenase (*SlNCED1*) [68,69], and an aldehyde oxidase [70], respectively. Both *flc* and *sit* are impaired in the conversion of ABA-aldehyde to ABA [67,70], while *not* is unable to catalyse the cleavage of 9-cis-violaxanthin and/or 9-cis-neoxanthin to xanthoxin, an ABA precursor [69]. Glasshouse-grown *flc*, *not* and *sit* mutants and control wild-type plants were subjected to water deprivation treatment and *Rider* transcript levels quantified by RT-qPCR (Figure 2D). *Rider* transcript levels were reduced in *flc*, *not* and *sit* by 43%, 26% and 56%, respectively.

To examine whether ABA stimulates accumulation of *Rider* transcripts, tomato seedlings were transferred to media supplemented with increasing concentrations of ABA (Figure 2E). The levels of *Rider* transcripts increased in a dose-dependent manner with increasing ABA concentrations. This suggests that ABA is not only involved in signalling that results in hyper-activation of *Rider* transcription during drought, but it also directly promotes the accumulation of *Rider* transcripts. The effectiveness of the treatments was verified by assaying expression of the stress- and ABA-responsive gene *SlASR1* (Figure S2A-F).

Identification in the U3 region of *Rider* LTRs of a binding domain for C-repeat binding factors (CBF), which are regulators of the cold-induced transcriptional cascade [64,71], led us to test *Rider* activation by cold stress. However, *Rider* transcription was not affected by cold treatment, leaving drought and salinity as the predominant environmental stresses identified so far that stimulate accumulation of *Rider* transcripts (Figure S2G).

### RdDM regulates levels of *Rider* transcripts

The suppression of transposon-derived transcription by epigenetic mechanisms, which typically include DNA methylation, maintains genome integrity [2,3,5]. We asked whether *Rider* transcription is also restricted by DNA methylation. Tomato seedlings were grown on media supplemented with 5-azacytidine, an inhibitor of DNA methyltransferases. *Rider* transcript steady-state levels increased in plants treated with 5-azacytidine compared to controls (Figure 3A). Comparison of *Rider* transcript accumulation in 5- azacytidine-treated and ABA-treated plants revealed similar levels of transcripts and the levels were similar when the treatments were applied together (*P* <0.05; Figure 3A).

**Figure 3:**
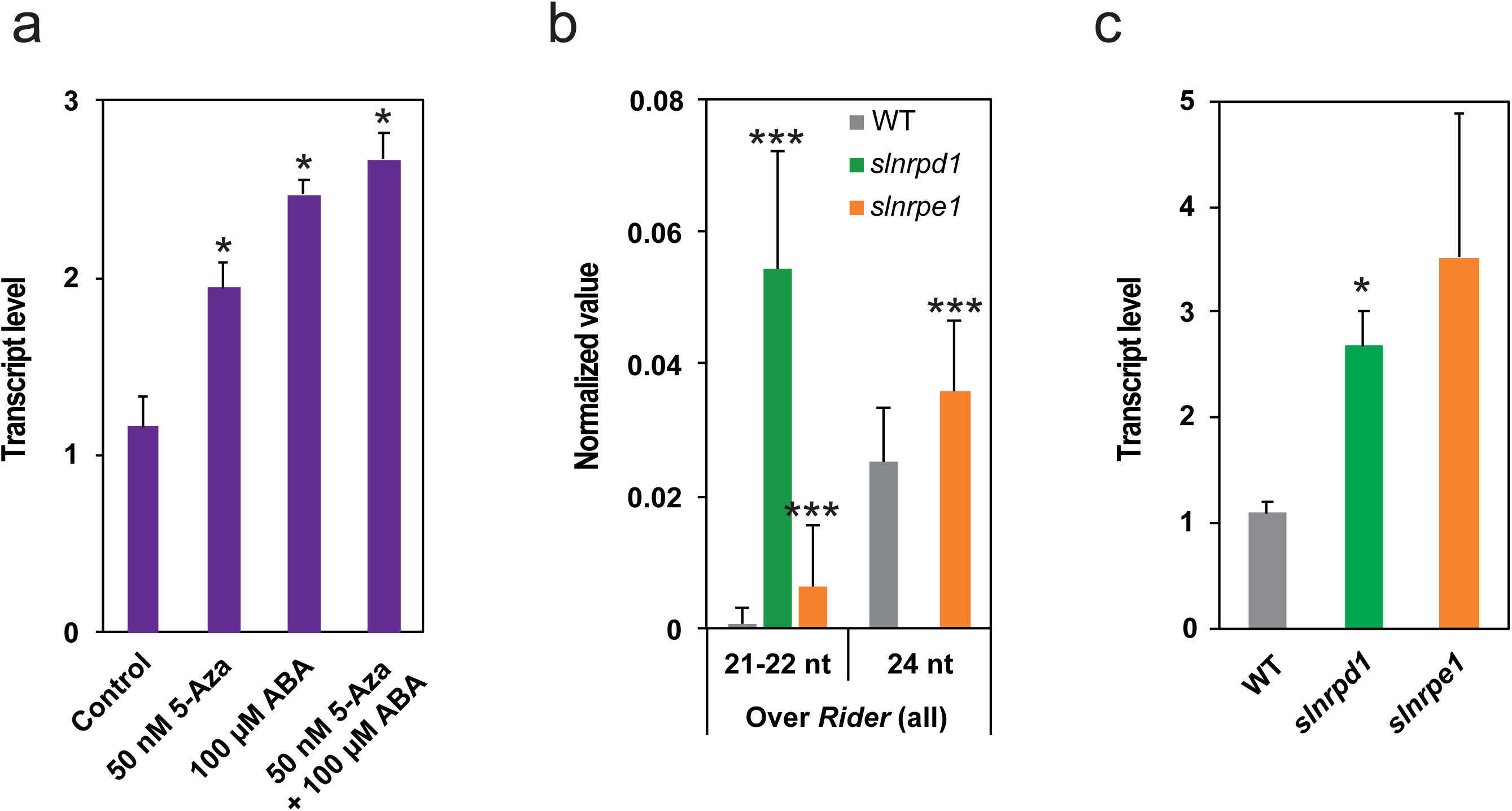
Accumulation of *Rider* transcripts in tomato plants deficient in epigenetic regulation. (A) Quantification of *Rider* RNA levels by RT-qPCR in tomato seedlings treated with 5-azacytidine and/or ABA. Histograms show normalized expression relative to Control, +/- SEM from two to three biological replicates. \**P*<0.05, two-sided Student’s *t*-test. (B) Abundance of siRNAs at *Rider* elements in wild type, *slnrpd1* and *slnrpe1*. Data are expressed as siRNA reads per kb per million mapped reads and represent average normalized siRNA counts on *Rider* elements +/- SD from 71 *de novo* annotated *Rider* copies. \*\*\**P*<0.001, two-sided Student’s *t*-test. (C) Quantification of *Rider* RNA by RT-qPCR in *slnrpd1* and *slnrpe1*. Histograms show normalized expression relative to WT, +/- SEM from two to four biological replicates. \**P*<0.05, two-sided Student’s *t*-test.

To further examine the role of DNA methylation in controlling *Rider* transcription, we took advantage of tomato mutants defective in crucial components of the RdDM pathway, namely SlNRPD1 and SlNRPE1, the major subunits of RNA Pol IV and Pol V, respectively. These mutants exhibit reduced cytosine methylation at CHG and CHH sites (in which H is any base other than G) residing mostly at the chromosome arms, with *slnrpd1* showing a dramatic, genome-wide loss of 24-nt siRNAs [47]. To evaluate the role of RdDM in *Rider* transcript accumulation, we first assessed the consequences of impaired RdDM on siRNA populations at full-length *Rider* elements. Deficiency in SlNRPD1 resulted in a complete loss of 24-nt siRNAs that target *Rider* elements (Figure 3B). This loss was accompanied by a dramatic increase (approximately 80-fold) in 21-22-nt siRNAs at *Rider* loci (Figure 3B). In contrast, the mutation in SlNRPE1 triggered increases in both 21-22-nt and 24-nt siRNAs targeting *Rider* elements (Figure 3B). We then asked whether altered distribution of these siRNA classes is related to the age of the *Rider* elements and/or their chromosomal position, and thus local chromatin properties. Compilation of the genomic positions and siRNA data in RdDM mutants didn’t reveal preferential accumulation of 21-22-nt siRNAs (Figure S3A) or 24-nt siRNAs (Figure S3B) over specific *Rider* classes. Subsequently, we examined whether loss of SlNRPD1 or SlNRPE1 was sufficient to increase levels of *Rider* transcripts and observed increased accumulation of *Rider* transcripts in both *slnrpd1* and *slnrpe1* compared to WT (Figure 3C).

We assessed whether this increase in *Rider* transcript levels is linked to changes in DNA methylation levels in *Rider* elements of RdDM mutants. There was no significant change in global DNA methylation in the three sequence contexts in the 71 *de novo* annotated *Rider* elements (Figure S3C), despite a tendency for young *Rider* elements to lose CHH in *slnrpd1* and *slnrpe1* (Figure S3D). Thus, the RdDM pathway affects the levels of *Rider* transcripts but there was no direct link to DNA methylation levels.

### Extrachromosomal circular DNA of *Rider* accumulates during drought stress and in *slnrpd1* and *slnrpe1* mutants

The life cycle of LTR retrotransposons starts with transcription of the element, then the synthesis and maturation of accessory proteins including reverse transcriptase and integrase, reverse transcription, and the production of extrachromosomal linear (ecl) DNA that integrates into a new genomic location [72]. In addition, eclDNA can be a target of DNA repair and can be circularised by a non-homologous end-joining mechanism or homologous recombination between LTRs, resulting in extrachromosomal circular DNA (eccDNA) [73–76]. We searched for eccDNA to evaluate the consequences of increased *Rider* transcript accumulation due to drought stress or an impaired RdDM pathway on subsequent steps of the transposition cycle. After exonuclease-mediated elimination of linear dsDNA and circular ssDNA, *Rider* eccDNA was amplified by sequence-specific inverse PCR (Figure 4A). *Rider* eccDNA was absent in plants grown in control conditions but was detected in plants subjected to drought stress (Figure 4A). Sanger sequencing of the inverse PCR products showed that the amplified eccDNA probably originates from the *Rider_08_3* copy, which has 98.2 % sequence homology of the 5’ and 3’ LTR sequences (Figure S4A). Residual sequence divergence may be due to genotypic differences between the reference genomic sequence and the genome of our experimental material. Analysis of CREs in the LTR of the eccDNA revealed the presence of all elements identified previously with the exception of a single nucleotide mutation located in the *CGCGBOXAT* box (Figure S4A). This suggests that while this CRE is not required for production of *Rider* eccDNA upon drought stress, presence of all other CREs including the two *MYBCORE* elements is likely to be necessary for its activation.

**Figure 4:**
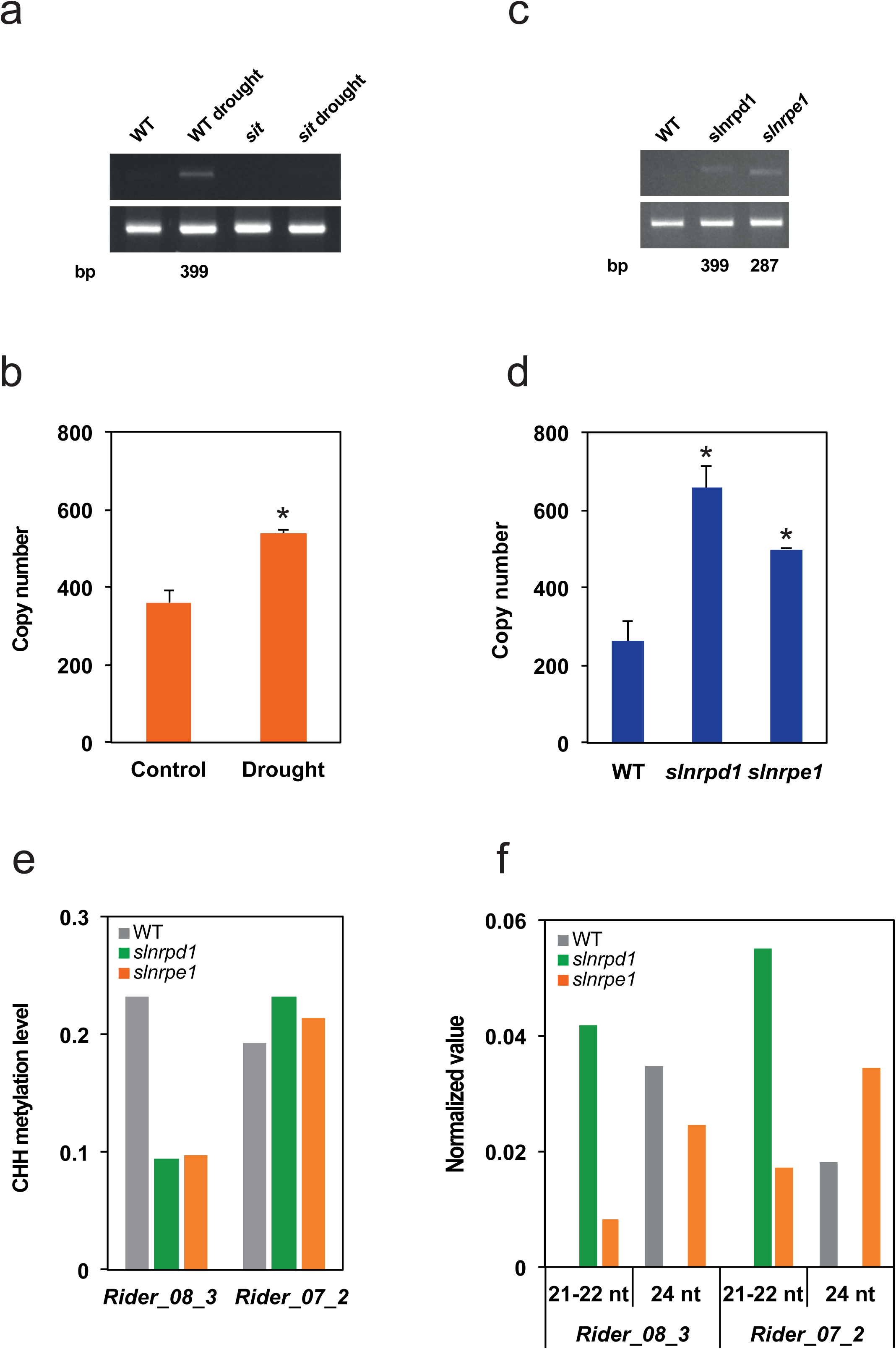
Accumulation of *Rider* extrachromosomal DNA in drought-stressed plants and in *slnrpd1* and *slnrpe1* mutants. (A) Assay by inverse PCR of *Rider* extrachromosomal circular DNA in drought-stressed wild-type plants and stressed *sit* mutants. Primer localization shown on the left (grey bar: *Rider* element, black box: LTR, arrowheads: PCR primers). Upper gel: specific PCR amplification of *Rider* circles after DNase treatment, lower gel: control PCR for *Rider* detection without DNase treatment. (B) Quantification of *Rider* DNA copy number, including both chromosomal and extrachromosomal copies, by qPCR in leaves of tomato plants subjected to drought-stress. Histograms show normalized expression +/- SEM from two to three biological replicates. \**P*<0.05, two-sided Student’s *t*-test. (C) Assay by inverse PCR of *Rider* extrachromosomal circular DNA in *slnrpd1* and *slnrpe1* leaves. Upper gel: PCR amplification of *Rider* circles after DNase treatment, lower gel: control PCR for *Rider* detection without DNase treatment. (D) Quantification of *Rider* DNA copy number, including both chromosomal and extrachromosomal copies, by qPCR in *slnrpd1* and *slnrpe1* leaves. Histograms show normalized expression +/- SEM from two biological replicates. \**P*<0.05, two-sided Student’s *t*-test. (E) Quantification of CHH DNA methylation levels at *Rider_08_3* and *Rider_07_2* in wild type, *slnrpd1* and *slnrpe1*. Levels expressed as % of methylated CHH sites. (F) Normalized siRNA count of 21-22-nt and 24-nt siRNAs at *Rider_08_3* and *Rider_07_2* in wild type, *slnrpd1* and *slnrpe1*. Data are expressed as siRNA reads per kb per million mapped reads.

Examination by quantitative PCR of the accumulation of *Rider* DNA, which included extrachromosomal and genomic copies, in drought-stressed plants also revealed an increase in *Rider* copy number due to eccDNA (Figure 4B). Importantly, *Rider* eccDNA was not detected in *sit* mutants subjected to drought stress (Figure 4A), suggesting that induced transcription of *Rider* by drought stress triggers production of extrachromosomal DNA and this response requires ABA biosynthesis.

We also examined the accumulation of *Rider* eccDNA in plants impaired in RdDM. Interestingly, *Rider* eccDNA was detected in *slnrpd1* and *slnrpe1* (Figure 4C) and increase in *Rider* DNA copy number due to eccDNA accumulation was confirmed by qPCR (Figure 4D). Absence of newly integrated genomic copies has been further validated by genome sequencing. The eccDNA forms differed between the mutants (Figure 4C). Sequencing of *Rider* eccDNA in *slnrpd1* showed a sequence identical to the *Rider* eccDNA of wild-type plants subjected to drought stress. Thus the *Rider_08_3* copy is probably the main contributor to eccDNA in drought and in *slnrpd1*. In contrast, eccDNA recovered from *slnrpe1* had a shorter LTR (287 bp) and the highest sequence similarity with *Rider_07_2* (89.2 %) (Figure S4B). Shortening of the LTR in this particular element is associated with the loss of the upstream *MYBCORE* as well as the *CGCGBOXAT* elements (Figure S4B). This suggests that in the absence of SlNRPE1, presence of these CREs is facultative for eccDNA production originating from this copy. In contrast, the absence of eccDNA copies derived from this element upon dehydration suggests that both *MYBCORE* elements are required for effective *Rider* activation upon drought stress.

We then asked whether DNA methylation and siRNA distribution at these particular *Rider* copies had changed in the mutants. DNA methylation at CHH sites, but not CG nor CHG, was drastically reduced at *Rider_08_3* in *slnrpd1* (Figure 4E and Figure S4C-E) together with a complete loss of 24-nt siRNAs at this locus (Figure 4F and Figure S4F) but DNA methylation at *Rider_07_2* was not affected, despite the deficiency of SlNRPD1 or SlNRPE1 (Figure 4E and Figure S4C-E). Levels of 21-22-nt siRNAs in both mutants and 24-nt siRNA in *slnrpe1* were increased (Figure 4F and Figure S4F-G). Altogether, this suggests that RdDM activity on *Rider* is highly copy-specific and that different components of the RdDM pathway differ in their effects on *Rider* silencing.

### *Rider* families in other plant species

To examine the distribution of *Rider* retrotransposons in other plant species, we searched for *Rider*-related sequences across the genomes of further *Solanaceae* species, including wild tomatoes, potato (*Solanum tuberosum*), and pepper (*Capsicum annuum*). We used the *Rider* reference sequence [44] as the query against genome sequences of *Solanum arcanum*, *S. habrochaites*, *S. lycopersicum*, *S. pennellii*, *S. pimpinellifolium*, *S. tuberosum*, and *Capsicum annuum* (genome versions are listed in Materials and Methods). Two BLAST searches were performed, one using the entire *Rider* sequence as the query and the other using only the *Rider* LTR. Consistent with previous reports, *Rider-like* elements are present in wild relatives of tomato such as *S. arcanum*, *S. pennellii* and *S. habrochaites;* however, the homology levels and their lengths vary significantly between species (Figure 5A). While *S. arcanum* and *S. habrochaites* exhibit high peak densities at 55% and 61% homology, respectively, *S. pennellii* show a high peak density at 98% over the entire *Rider* reference sequence (Figure 5A). This suggests that the *S. arcanum* and *S. habrochaites* genomes harbour mostly *Rider-like* elements with relatively low sequence similarity, while *S. pennellii* retains full-length *Rider* elements.

**Figure 5:**
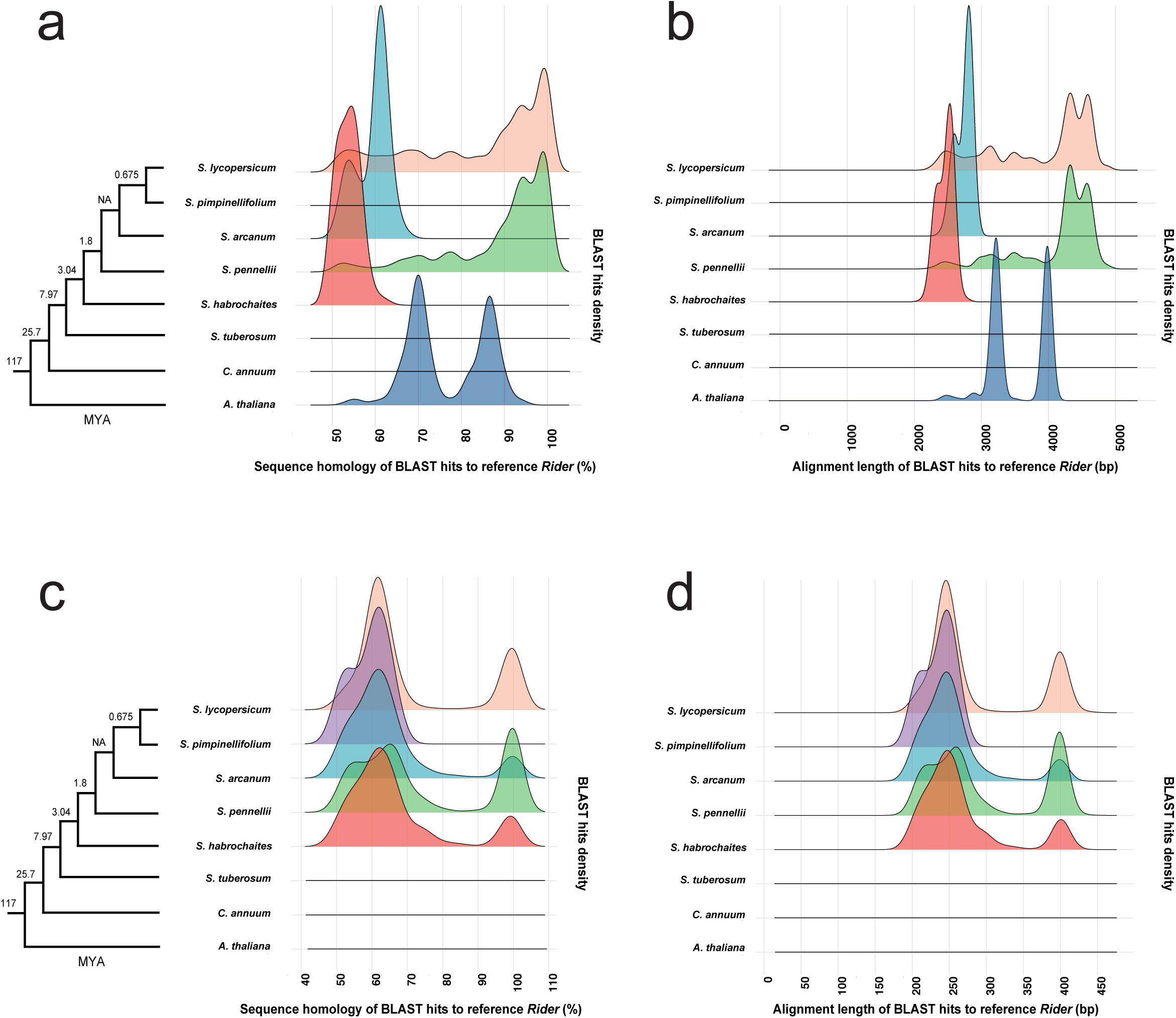
Distribution of *Rider* in other *Solanaceae* species. (A) *In silico* identification of *Rider* elements in *Solanaceae* species based on the density of high homology BLAST hits over the full-length reference *Rider* sequence. (B) Alignment length of high homology BLAST hits obtained in (A). (C) *In silico* identification of *Rider* elements in *Solanaceae* species based on the density of high homology BLAST hits over the reference *Rider* LTR sequence. (D) Alignment length of high homology BLAST hits obtained in (C). Left panels (A) and (C): phylogenetic trees of the species examined.

To better visualize this situation, we aligned the BLAST hits to the reference *Rider* copy (Figure 5B). This confirmed that *Rider* elements in *S. pennellii* are indeed mostly full-length *Rider* homologs showing high density of hits throughout their lengths, while BLAST hits in the *S. arcanum* and *S. habrochaites* genomes showed only partial matches over the 4867 bp of the reference *Rider* sequence (Figure 5B). Unexpectedly, this approach failed to detect either full-length or truncated *Rider* homologs in the close relative *of* tomato, *S. pimpinellifolium*. Extension of the same approaches to the genomes of the evolutionary more distant *S. tuberosum* and *Capsicum annuum* failed to detect substantial *Rider* homologs (Figure 5A-B), confirming the absence of *Rider* in the potato and pepper genomes [44]. As a control, we also analysed *Arabidopsis thaliana*, since previous studies reported the presence of *Rider* homologs in this model plant [44]. Using the BLAST approach above, we repeated the results provided in [44] and found BLAST hits of high sequence homology to internal sequences of *Rider* in the *Arabidopsis thaliana* genome. However, we did not detect sequence homologies to *Rider* LTRs (Figure 5C-D). Motivated by this finding and the possibility that *Rider* homologs in other species may have highly divergent LTRs, we screened for *Rider* LTRs that would have been missed in the analysis shown in Figure 5A-B due to the use of the full-length sequence of *Rider* as the query. Using the *Rider* LTR as a query revealed that *S. pennellii, S. arcanum* and *S. habrochaites* retain intact *Rider* LTR homologs, but *S. pimpinellifolium* exhibits a high BLAST hit density exclusively at approximately 60% homology. This suggests strong divergence of *Rider* LTRs in this species (Figure 5C-D). Overall, the results indicate intact *Rider* homologs in some *Solanaceae* species, whereas sequence similarities to *Rider* occur only within the coding area of the retrotransposons in more distant plants such as *Arabidopsis thaliana.* Therefore, LTRs, which include the *cis*-regulatory elements conferring stress-responsiveness, diverge markedly between species.

To address the specificity of this divergence in *Solanaceae* species, we examined whether the CREs enriched in *S. lycopersicum* (Figure 2A) are present in LTR sequences of the *Rider* elements in *S. pennellii, S. arcanum*, *S. habrochaites* and *S. pimpinellifolium* (Figure 5C). While the LTRs identified in *S. pennellii, S. arcanum* and *S. habrochaites* retained all five previously identified CREs, more distant LTRs showed shortening of the U3 region associated with loss of the CGCG box (Figure S5 and Table S4). This was observed already in *S. pimpinellifolium*, where all identified *Rider* LTRs lacked part of the U3 region containing the CGCG box (Supplementary Figure 5). Thus, *Rider* distribution and associated features differ even between closely related *Solanaceae* species, correlated with the occurrence of a truncated U3 region and family-wide loss of CREs.

Finally, to test the evolutionary conservation of *Rider* elements across the plant kingdom, we performed *Rider* BLAST searches against all 110 plant genomes available at the NCBI Reference Sequence (RefSeq) database (www.ncbi.nlm.nih.gov/refseq). Using the entire *Rider* sequence as the query to measure the abundance of *Rider* homologs throughout these genomes, we found *Rider* homologs in 14 diverse plant species (Figure S6). This suggests that *Rider* in tomato did not originate by horizontal transfer from Arabidopsis as initially suggested [44], but rather that *Rider* was already present in the last common ancestor of these plant species and persisted or was subsequently eradicated from the genomes. The limited conservation of *Rider* LTR sequences in the same 14 species, revealed using the LTR sequence as the query, suggests that *Rider* LTRs are rapidly evolving and that drought-responsive CREs may be restricted to *Solanaceae* (Figure S7).

## DISCUSSION

### High-resolution map of full-length Rider elements in the tomato genome

Comprehensive analysis of individual LTR retrotransposon families in complex plant genomes has been facilitated and become more accurate with the increasing availability of high-quality genome assemblies. Here, we took advantage of the most recent tomato genome release (SL3.0) to characterize with improved resolution the high-copy-number *Rider* retrotransposon family. Although *Rider* activity has been causally linked to the emergence of important agronomic phenotypes in tomato, the triggers of *Rider* have remained elusive. Despite the relatively low proportion (approximately 20%) of euchromatic chromosomal regions in the tomato genome [31]), our *de novo* functional annotation of full-length *Rider* elements revealed preferential compartmentalization of recent *Rider* insertions within euchromatin compared to aged insertions. Mapping analyses further revealed that recent rather than aged *Rider* transposition events are more likely to modify the close vicinity of genes. However, *Rider* copies inserted into heterochromatin have been passively maintained for longer periods. This differs significantly from other retrotransposon families in tomato such as *Tnt1*, *ToRTL1* and *T135*, which show initial, preferential insertions into heterochromatic regions [77]. *TARE1*, a high-copy-number *Copia-like* element, is present predominantly in pericentromeric heterochromatin [78]. Another high-copy-number retrotransposon, *Jinling*, is also enriched in heterochromatic regions, making up about 2.5% of the tomato nuclear genome [79]. The *Rider* propensity to insert into gene-rich areas mirrors the insertional preferences of the *ONSEN* family in Arabidopsis. Since new *ONSEN* insertions confer heat-responsiveness to neighbouring genes [28,29], it is tempting to speculate that genes in the vicinity of new *Rider* insertions may acquire, at least transiently, drought-responsiveness.

### Environmental and epigenetic regulation of Rider activity

We found that *Rider* transcript levels are elevated during dehydration stress mediated by ABA-dependent signalling. The activation of retrotransposons upon environmental cues has been shown extensively to rely on the presence of *cis*-regulatory elements within the retrotransposon LTRs [60]. The heat-responsiveness of *ONSEN* in Arabidopsis [26,27,80], *Go-on* in rice [25], and *Copia* in Drosophila [81] is conferred by the presence in their LTRs of consensus sequences found in the promoters of heat-shock responsive genes. Thus, the host’s heat-stress signalling appears to induce transcriptional activation of the transposon and promote transposition [80]. While *ONSEN* and *Go-on* are transcriptionally inert in the absence of a triggering stress, transcripts of Drosophila *Copia* are found in control conditions, resembling the regulatory situation in *Rider*. Due to relatively high constitutive expression, increase in transcript levels of Drosophila *Copia* following stress appears modest compared to *ONSEN* or *Go-on*, which are virtually silent in control conditions [25–27,80]. Regulation of Drosophila *Copia* mirrors that of *Rider*, where transcript levels during dehydration stress are very high but the relative increase compared to control conditions is rather modest.

The presence of MYB recognition sequences within *Rider* LTRs suggests that MYB transcription factors participate in transcriptional activation of *Rider* during dehydration. Multiple MYB subfamilies are involved in ABA-dependent stress responses in tomato, but strong enrichment of the MYB core element CTGTTA within *Rider* LTRs suggests involvement of R2R3-MYB transcription factors, which are markedly amplified in *Solanaceae* [82]. Members of this MYB subfamily are involved in the ABA signalling-mediated drought-stress response [83] and salt-stress signalling [84]. This possible involvement of R2R3-MYBs in *Rider* is reminiscent of the transcriptional activation of the tobacco retrotransposon *Tto1* by the R2R3-MYB, member NtMYB2 [85]. Drought-responsiveness has been observed for *Rider_08_3* only, despite other individual *Rider* copies displaying intact MYB core element (Table S7). This suggests that presence of this CRE is not the only feature required for drought-responsiveness, and other factors, such as genomic location, influence *Rider* activity.

In addition to environmental triggers, *Rider* transcript levels are regulated by the RdDM pathway. Depletion of SlNRPD1 and SlNRPE1 increases *Rider* transcript abundance, resulting in production of extrachromosomal circular DNA. Analysis of *Rider*-specific siRNA populations revealed that siRNA targeting of *Rider* elements is mostly independent of their genomic location and chromatin context. This is somewhat unexpected since RdDM activity in tomato seems to be restricted to gene-rich euchromatin and it was postulated that accessibility of RNA Pol IV to heterochromatin is hindered by the compact chromatin structure [47,86,87]. We identified *Rider* copies targeted by RdDM, which potentially influences local epigenetic features. Loss of SlNRPD1 and SlNRPE1 leads to over-accumulation of 21- 22-nt siRNAs at *Rider* copies, suggesting that inactivation of canonical RdDM pathway-dependent transcriptional gene silencing triggers the activity of the non-canonical RDR6 RdDM pathway at *Rider* [88–90].

It is noteworthy that, despite clear effects on *Rider* transcript accumulation and siRNA accumulation, loss of SlNRPD1 and SlNRPE1 is not manifested by drastic changes in total DNA methylation levels of *Rider* at the family level. This is in accordance with the modest decrease in genome-wide CHH and CHG methylation described in tomato RdDM mutants, with most of the changes happening on the euchromatic arms while the pericentromeric heterochromatin is unaffected [47]. Distribution of the 71 intact *Rider* elements in both euchromatic and heterochromatic compartments thus likely hampers detection of major changes DNA methylation over the *Rider* family. Only young euchromatic *Rider* elements marginally lose CHH methylation in the *slnrpd1* mutant, but this is modest compared to the general decrease in mCHH observed throughout the chromosome arms [47]. As expected, CHH methylation at heterochromatic *Rider* is not affected. This suggests that SlCMT2 is involved in maintenance of mCHH at heterochromatic *Rider* copies in the absence of SlNRPD1, as observed previously for pericentromeric heterochromatin [47]. In general, our observations suggest that epigenetic silencing of *Rider* retrotransposons is particularly robust and involves compensatory pathways.

We identified extrachromosomal circular DNA originating from the *Rider* copies *Rider_08_3* and *Rider_07_2* in *slnrpd1* and *slnrpe1*, respectively. In terms of DNA methylation and siRNA distribution at these two specific copies, loss of SlNRPD1 and SlNRPE1 brought different copy-specific outcomes. *Rider_08_3*, the main contributor to eccDNA in *slnrpd1*, displayed a reduction in CHH methylation that may contribute to increased transcription and the accumulation of eccDNA. In *Rider_07_2*, that provides a template for eccDNA in *slnrpe1*, there was no change in DNA methylation levels. Therefore, transcription and the production of eccDNA from this *Rider* copy is not regulated by DNA methylation. Consequently, eccDNA from *Rider_07_2* was not detected in *slnrpd1* despite drastic loss of CHH methylation.

Despite our efforts, we were unable to apply either drought or ABA treatment to the *slnrpd1* and *slnrpe1* mutants. In contrast to Arabidopsis [91,92], RdDM mutants in tomato are showing severe developmental defects and are sterile [47]. They are particularly difficult to maintain, precluding the application of stress treatments. Altogether, it appears that transcriptional control and reverse transcription of *Rider* copies occurs via multiple layers of regulation, possibly specific for individual *Rider* elements according to age, sequence or genomic location, that are targeted by parallel silencing pathways, including non-canonical RdDM [93,94].

### Rider retrotransposons in other plant species

The presence of *Rider* in tomato relatives as well as in more distantly related plant species has been described previously [33,44,46]. However, the *de novo* identification of *Rider* elements in the sampling provided here shows the distribution of the *Rider* family within plant species to be more complex than initially suggested. Surprisingly, mining for sequences with high similarity, overlapping more than 85% of the entire reference sequence of *Rider*, detected no full-length *Rider* elements in *Solanum pimpinellifolium* but in all other wild tomato species tested. Furthermore, the significant accumulation of only partial *Rider* copies in *Solanum pimpinellifolium*, the closest relative of tomato, does not match the established phylogeny of the *Solanaceae*. The cause of these patterns is unresolved but two scenarios can be envisaged. First, the absence of full-length *Rider* elements may be due to the suboptimal quality of genome assembly that may exclude a significant proportion of highly repetitive sequences such as *Rider*. This is supported by the N50 values within the *Solanaceae*, where the quality of genome assemblies varies significantly between species, with *S. pimpinellifolium* showing the lowest (Table S8). An improved genome assembly would allow a refined analysis of *Rider* in this species. Alternatively, active *Rider* copies may have been lost in *S. pimpinellifolium* since the separation from the last common ancestor but not in the *S. lycopersicum* and *S. pennellii* lineages. The high-density of solo-LTRs and truncated elements in *S. pimpinellifolium* is in agreement with this hypothesis.

Comparing the sequences of *Rider* LTRs in the five tomato species, the unique occurrence of LTRs lacking most of the U3 region in *S. pimpinnellifolium* suggests that loss of important regulatory sequences has impeded maintenance of intact *Rider* elements. Interestingly, part of the U3 region missing in *S. pimpinellifolium* contains the CGCG box, which is involved in response to environmental signals [61], as well as a short CpG-island-like structure (position 52-155 bp on reference *Rider*). CpG islands are usually enriched 5’ of transcriptionally active genes in vertebrates [95] and plants [96]. Despite the presence of truncated *Rider* LTRs, the occurrence of intact, full-length LTRs in other wild tomato species indicates that *Rider* is still potentially active in these genomes.

Altogether, our findings suggest that inter- and intra-species TE distribution can be uncoupled and that the evolution of TE families in present crop plants was more complex than initially anticipated. Finally, we have opened interesting perspectives for harnessing transposon activities in crop breeding. Potentially active TE families that react to environmental stimuli, such as *Rider*, provide an unprecedented opportunity to generate genetic and epigenetic variation from which desirable agronomical traits may emerge. Notably, rewiring of gene expression networks regulating the drought-stress responses of new *Rider* insertions is an interesting strategy to engineer drought-resilient crops.

## DATA AVAILABILITY

SRAtoolkit, v2.8.0 (https://github.com/ncbi/sra-tools) and Biomartr 0.9.9000 (https://ropensci.github.io/biomartr/index.html) were used for data collection.

Phylogenetic trees were constructed using Geneious 9.1.8 (www.geneious.com).

The *de novo* retrotransposon annotation pipeline *LTRpred* is available in the GitHub repository (https://github.com/HajkD/LTRpred).

*Rider* annotation and analysis pipeline is available in the GitHub repository (https://github.com/HajkD/RIDER).

Distribution of *Rider* elements was done using the R package *metablastr* (https://github.com/HajkD/metablastr).

DNA methylation levels were assessed using the R package DMRcaller (http://bioconductor.org/packages/release/bioc/html/DMRcaller.html).

Small RNA analysis was done using Trim Galore! (www.bioinformatics.babraham.ac.uk/projects/trim_galore), ShortStack v3.6 (https://github.com/MikeAxtell/ShortStack) and GenomicRanges v3.8 (https://bioconductor.org/packages/release/bioc/html/GenomicRanges.html).

Reference *Rider* nucleotide sequence (accession number EU195798) is available here (https://www.ncbi.nlm.nih.gov/nuccore/EU195798).

Public sequencing data used in this study are available at Sequence Read Archive (SRA) (https://www.ncbi.nlm.nih.gov/sra/) under accession numbers “SRP081115”, “SRR4013319”, “SRR4013316”, “SRR4013314” and “SRR4013312”.

## ACKNOWLEDGMENTS

The authors thank all members of Dr. Paszkowski lab for fruitful discussions during the development of this project as well as the SLCU support staff.

## Author contributions

MB and JP designed the study; MB performed experiments; MB, HGD and MC performed genomic data analyses; QG, SLG and DCB provided unpublished material and data; MB and JP wrote the paper with contributions from HGD and MC. All authors read and approved the final manuscript.

## SUPPLEMENTARY FIGURE LEGENDS

**Figure S1: Distribution of 71 *de novo* annotated *Rider* elements based on LTR similarity and chromatin context**

(A) Age distribution of total *Rider* elements based on LTR similarity and corresponding classes. (B) Age distribution of *Rider* elements inserted in heterochromatic (HC) and euchromatic (EC) regions based on LTR similarity.

**Figure S2: *Rider* transcripts levels are unaffected by cold stress**

(A-D) Quantification of *SlASR1* RNA levels by RT-qPCR in wild-type tomato seedlings after (A) drought stress (B) mannitol, (C) NaCl or (D) ABA treatments. (E) Quantification of *SlASR1* RNA levels in leaves of drought-stressed tomato wild-type plants, *flc*, *not* and *sit* mutants. (F) Quantification of *SlASR1* RNA levels by RT-qPCR in wild-type tomato seedlings after 5- azacytidine and ABA treatments. (G) Quantification of *Rider* RNA levels by RT-qPCR in wild-type tomato seedlings after cold stress. Histograms show normalized expression +/- SEM from three to five biological replicates.

**Figure S3: Distribution of siRNAs and DNA methylation within *Rider* sub-groups**

(A) 21-22-nt and (B) 24-nt siRNAs normalized counts at distinct *Rider* sub-groups in wild type, wild type with *CAS9*, *slnrpd1* and *slnrpe1*. *Rider* elements are classified based on LTR similarity (80-95%, 95-98% and 98-100%), while *Rider* (Euchromatin) denotes copies located on euchromatic arms and *Rider* (Heterochromatin) copies located in pericentromeric heterochromatin. Data are expressed as siRNA reads per kb per million mapped reads, and represent average normalized siRNA counts on *Rider* elements +/- SD from *Rider* copies in the sub-group. (C) Quantification of DNA methylation levels in the CG, CHG and CHH contexts at *Rider* in wild type, *slnrpd1* and *slnrpe1*. The levels are averages of DNA methylation (%) in each context over the 71 *de novo* annotated *Rider* copies. (D) Quantification of CHH DNA methylation levels at *Rider* sub-groups in wild type, *slnrpd1* and *slnrpe1*. The levels are averages of DNA methylation (%) in the CHH context over *Rider* sub-groups.

**Figure S4: Distinct *Rider* copies contribute to the production of extrachromosomal circular DNA**

Comparison of the LTR nucleotide sequence from *Rider* extrachromosomal circular DNA detected after drought, or in *slnrpd1* (A) or *slnrpe1* (B), with the reference *Rider* LTR using EMBOSS Needle (www.ebi.ac.uk/Tools/psa/emboss_needle). CREs are marked as coloured boxes. (C) Quantification of CHH DNA methylation levels at LTRs and body of *Rider_08_3* and *Rider_07_2* in wild type, *slnrpd1* and *slnrpe1*. Levels expressed as % of methylated CHH sites. (D-E) Quantification of CG (D) and CHG (E) DNA methylation levels at *Rider_08_3* and *Rider_07_2* in wild type, *slnrpd1* and *slnrpe1*. Levels expressed as % of methylated sites. (F-G) Normalized siRNA count of 24-nt (F) and 21-22-nt (G) siRNAs at LTRs and body of *Rider_08_3* and *Rider_07_2* in wild type, *slnrpd1* and *slnrpe1*. Data are expressed as siRNA reads per kb per million mapped reads.

**Figure S5: Characterization of *Rider* sub-populations in *Solanaceae* based on LTR sequences**

Coverage over reference *Rider* LTR of high homology sequences identified by BLAST in Figure 5C. Sequences classified as “long LTR” were selected by filtering for BLAST hits with alignment lengths between 350-450 bp and >50% sequence and length homology to reference *Rider*. Sequences classified as “short LTR” were selected by filtering for BLAST hits with alignment lengths between 150-300 bp and >50% sequence and length homology to reference *Rider*.

**Figure S6: Identification of *Rider* homologs in 14 plant species**

*In silico* identification of *Rider* homologs in 14 plant species based on the density of high homology BLAST hits over the full-length reference *Rider* sequence (left) and alignment length of BLAST hits obtained (right). Species are ordered by evolutionary distance to *Solanum lycopersicum* from www.timetree.org, www.genome.jp and Supplementary References.

**Figure S7: Non-*Solanaceae Rider* homologs lack LTR sequence conservation**

*In silico* identification of *Rider* LTR homologs in 14 plant species based on the density of high homology BLAST hits over the reference *Rider* LTR sequence only. Species are ordered by evolutionary distance to *Solanum lycopersicum* from www.timetree.org, www.genome.jp and Supplementary References.

## SUPPLEMENTARY TABLES LEGENDS

**Table S1: Primers used in this study**

**Table S2: List of the 110 plant species used for the large-scale *Rider* BLAST search**

**Table S3: Identification and enrichment analysis of *cis*-regulatory elements in *Rider* LTRs**

**Table S4: Enrichment analysis of *cis*-regulatory elements in *Rider* LTRs in four *Solanaceae* species**

**Table S5: *De novo* annotation of LTR retrotransposons in the SL3.0 genome by *LTRpred***

**Table S6: Patristic distances between 71 *de novo* annotated *Rider* copies**

**Table S7: Presence of *cis*-regulatory elements in individual *Rider* copies**

**Table S8: N50 metric for six *Solanaceae* species**

## REFERENCES

1. Lisch D (2013) How important are transposons for plant evolution? Nat Rev Genet 14: 49–61.

2. Lisch D (2009) Epigenetic Regulation of Transposable Elements in Plants. Annu Rev Plant Biol 60: 43–66.

3. Slotkin RK, Martienssen R (2007) Transposable elements and the epigenetic regulation of the genome. Nat Rev Genet 8: 272–285.

4. Zhang H, Zhu J-K (2011) RNA-directed DNA methylation. Curr Opin Plant Biol 14: 142–147.

5. Rigal M, Mathieu O (2011) A ‘mille-feuille’ of silencing: Epigenetic control of transposable elements. Biochim Biophys Acta - Gene Regul Mech 1809: 452–458.

6. Law JA, Jacobsen SE (2010) Establishing, maintaining and modifying DNA methylation patterns in plants and animals. Nat Rev Genet 11: 204–220.

7. Matzke M, Kanno T, Huettel B, Daxinger L, Matzke AJM (2007) Targets of RNA-directed DNA methylation. Curr Opin Plant Biol 10: 512–519.

8. Wendte JM, Pikaard CS (2017) The RNAs of RNA-directed DNA methylation. Biochim Biophys Acta 1860: 140–148.

9. Mirouze M, Reinders J, Bucher E, Nishimura T, Schneeberger K, Ossowski S, Cao J, Weigel D, Paszkowski J, Mathieu O (2009) Selective epigenetic control of retrotransposition in Arabidopsis. Nature 461: 1–5.

10. Kato M, Miura A, Bender J, Jacobsen SE, Kakutani T (2003) Role of CG and non-CG methylation in immobilization of transposons in Arabidopsis. Curr Biol 13: 421–426.

11. Lanciano S, Carpentier MC, Llauro C, Jobet E, Robakowska-Hyzorek D, Lasserre E, Ghesquière A, Panaud O, Mirouze M (2017) Sequencing the extrachromosomal circular mobilome reveals retrotransposon activity in plants. PLoS Genet 13: 1–20.

12. Hu L, Li N, Xu C, Zhong S, Lin X, Yang J, Zhou T, Yuliang A, Wu Y, Chen Y-R, et al. (2014) Mutation of a major CG methylase in rice causes genome-wide hypomethylation, dysregulated genome expression, and seedling lethality. Proc Natl Acad Sci 111: 10642–10647.

13. Cheng C, Tarutani Y, Miyao A, Ito T, Yamazaki M, Sakai H, Fukai E, Hirochika H (2015) Loss of function mutations in the rice chromomethylase OsCMT3a cause a burst of transposition. Plant J 83: 1069–1081.

14. Miura A, Yonebayashi S, Watanabe K, Toyama T, Shimada H, Kakutani T (2001) Mobilization of transposons by a mutation abolishing full DNA methylation in Arabidopsis. Nature 411: 212–214.

15. Lippman Z, Gendrel A-V, Black M, Vaughn MW, Dedhia N, McCombie WR, Lavine K, Mittal V, May B, Kasschau KD, et al. (2004) Role of transposable elements in heterochromatin and epigenetic control. Nature 430: 471–476.

16. Tsukahara S, Kobayashi A, Kawabe A, Mathieu O, Miura A, Kakutani T (2009) Bursts of retrotransposition reproduced in Arabidopsis. Nature 461: 423–426.

17. Griffiths J, Catoni M, Iwasaki M, Paszkowski J (2018) Sequence-Independent Identification of Active LTR Retrotransposons in Arabidopsis. Mol Plant 11: 508–511.

18. Tan F, Zhou C, Zhou Q, Zhou S, Yang W, Zhao Y, Li G, Zhou D-X (2016) Analysis of Chromatin Regulators Reveals Specific Features of Rice DNA Methylation Pathways. Plant Physiol 171: 2041–2054.

19. Chuong EB, Elde NC, Feschotte C (2017) Regulatory activities of transposable elements: from conflicts to benefits. Nat Rev Genet 18: 71–86.

20. McClintock B (1951) Chromosome Organization and Genic Expression. Cold Spring Harb Symp Quant Biol 16: 13–47.

21. Grandbastien MA (1998) Activation of plant retrotransposons under stress conditions. Trends Plant Sci 3: 181–187.

22. Grandbastien MA, Audeon C, Bonnivard E, Casacuberta JM, Chalhoub B, Costa APP, Le QH, Melayah D, Petit M, Poncet C, et al. (2005) Stress activation and genomic impact of Tnt1 retrotransposons in Solanaceae. Cytogenet Genome Res 110: 229–241.

23. Butelli E, Licciardello C, Zhang Y, Liu J, Mackay S, Bailey P, Reforgiato-Recupero G, Martin C (2012) Retrotransposons Control Fruit-Specific, Cold-Dependent Accumulation of Anthocyanins in Blood Oranges. Plant Cell 24: 1242–1255.

24. Johns M a, Mottinger J, Freeling M (1985) A low copy number, copia-like transposon in maize. EMBO J 4: 1093–1101.

25. Cho J, Benoit M, Catoni M, Drost H-G, Brestovitsky A, Oosterbeek M, Paszkowski J (2018) Sensitive detection of pre-integration intermediates of long terminal repeat retrotransposons in crop plants. Nat Plants 317479.

26. Tittel-Elmer M, Bucher E, Broger L, Mathieu O, Paszkowski J, Vaillant I (2010) Stress-Induced Activation of Heterochromatic Transcription. PLoS Genet 6: e1001175.

27. Pecinka A, Dinh HQ, Baubec T, Rosa M, Lettner N, Scheid OM (2010) Epigenetic Regulation of Repetitive Elements Is Attenuated by Prolonged Heat Stress in *Arabidopsis*. Plant Cell 22: 3118–3129.

28. Ito H, Gaubert H, Bucher E, Mirouze M, Vaillant I, Paszkowski J (2011) An siRNA pathway prevents transgenerational retrotransposition in plants subjected to stress. Nature 472: 115–119.

29. Gaubert H, Sanchez DH, Drost H-G, Paszkowski J (2017) Developmental Restriction of Retrotransposition Activated in Arabidopsis by Environmental Stress. Genetics 207: 813–821.

30. Tenaillon MI, Hollister JD, Gaut BS (2010) A triptych of the evolution of plant transposable elements. Trends Plant Sci 15: 471–478.

31. Sato S, Tabata S, Hirakawa H, Asamizu E, Shirasawa K, Isobe S, Kaneko T, Nakamura Y, Shibata D, Aoki K, et al. (2012) The tomato genome sequence provides insights into fleshy fruit evolution. Nature 485: 635–641.

32. Xiao H, Jiang N, Schaffner E, Stockinger EJ, van der Knaap E (2008) A Retrotransposon-Mediated Gene Duplication Underlies Morphological Variation of Tomato Fruit. Science (80-) 319: 1527–1530.

33. Jiang N, Gao D, Xiao H, van der Knaap E (2009) Genome organization of the tomato sun locus and characterization of the unusual retrotransposon Rider. Plant J 60: 181–193.

34. Rodríguez GR, Muños S, Anderson C, Sim S-C, Michel A, Causse M, Gardener BBM, Francis D, van der Knaap E (2011) Distribution of SUN, OVATE, LC, and FAS in the tomato germplasm and the relationship to fruit shape diversity. Plant Physiol 156: 275–285.

35. Reynard GB (1961) New source of the j2 gene governing Jointless pedicel in tomato. Science (80-) 134: 2102.

36. Rick CM (1956) A new jointless gene from the Galapagos L. pimpinellifolium. TGC Rep 23.

37. Rick CM (1956) Genetic and Systematic Studies on Accessions of Lycospersicon from the Galapagos Islands. Am J Bot 43: 687.

38. Soyk S, Lemmon ZH, Oved M, Fisher J, Liberatore KL, Park SJ, Goren A, Jiang K, Ramos A, van der Knaap E, et al. (2017) Bypassing Negative Epistasis on Yield in Tomato Imposed by a Domestication Gene. Cell 169: 1142–1155.e12.

39. Fray RG, Grierson D (1993) Identification and genetic analysis of normal and mutant phytoene synthase genes of tomato by sequencing, complementation and co-suppression. Plant Mol Biol 22: 589–602.

40. Jiang N, Visa S, Wu S, Knaap E Van Der (2012) Rider Transposon Insertion and Phenotypic Change in Tomato. Springer Berlin Heidelberg, Berlin, Heidelberg.

41. Busch BL, Schmitz G, Rossmann S, Piron F, Ding J, Bendahmane A, Theres K (2011) Shoot Branching and Leaf Dissection in Tomato Are Regulated by Homologous Gene Modules. Plant Cell 23: 3595–3609.

42. Brown JC, Chaney RL, Ambler JE (1971) A New Tomato Mutant Inefficient in the Transport of Iron. Physiol Plant 25: 48–53.

43. Ling H-Q, Bauer P, Bereczky Z, Keller B, Ganal M (2002) The tomato fer gene encoding a bHLH protein controls iron-uptake responses in roots. Proc Natl Acad Sci U S A 99: 13938–13943.

44. Cheng X, Zhang D, Cheng Z, Keller B, Ling H-Q (2009) A New Family of Ty1-copia-Like Retrotransposons Originated in the Tomato Genome by a Recent Horizontal Transfer Event. Genetics 181: 1183–1193.

45. Wang Y, Diehl A, Wu F, Vrebalov J, Giovannoni J, Siepel A, Tanksley SD (2008) Sequencing and comparative analysis of a conserved syntenic segment in the Solanaceae. Genetics 180: 391–408.

46. Gilbert C, Feschotte C (2018) Horizontal acquisition of transposable elements and viral sequences: patterns and consequences. Curr Opin Genet Dev 49: 15–24.

47. Gouil Q, Baulcombe DC (2016) DNA Methylation Signatures of the Plant Chromomethyltransferases. PLoS Genet 12: 1–17.

48. Tanksley SD, Ganal MW, Prince JP, De Vicente MC, Bonierbale MW, Broun P, Fulton TM, Giovannoni JJ, Grandillo S, Martin GB, et al. (1992) High density molecular linkage maps of the tomato and potato genomes. Genetics 132: 1141–1160.

49. Catoni M, Griffiths J, Becker C, Zabet NR, Bayon C, Dapp M, Lieberman-Lazarovich M, Weigel D, Paszkowski J (2017) DNA sequence properties that predict susceptibility to epiallelic switching. EMBO J 36: 617–628.

50. Catoni M, Tsang JM, Greco AP, Zabet NR (2018) DMRcaller: a versatile R/Bioconductor package for detection and visualization of differentially methylated regions in CpG and non-CpG contexts. Nucleic Acids Res 1–11.

51. Lawrence M, Huber W, Pagès H, Aboyoun P, Carlson M, Gentleman R, Morgan MT, Carey VJ (2013) Software for Computing and Annotating Genomic Ranges. PLoS Comput Biol 9: 1–10.

52. Pruitt KD, Tatusova T, Maglott DR (2007) NCBI reference sequences (RefSeq): a curated non-redundant sequence database of genomes, transcripts and proteins. Nucleic Acids Res 35: D61–5.

53. Drost H-G, Paszkowski J (2017) Biomartr: genomic data retrieval with R. Bioinformatics 33: 1216–1217.

54. Fernandez-Pozo N, Menda N, Edwards JD, Saha S, Tecle IY, Strickler SR, Bombarely A, Fisher-York T, Pujar A, Foerster H, et al. (2015) The Sol Genomics Network (SGN)—from genotype to phenotype to breeding. Nucleic Acids Res 43: D1036–D1041.

55. Lowe TM, Eddy SR (1996) TRNAscan-SE: A program for improved detection of transfer RNA genes in genomic sequence. Nucleic Acids Res 25: 955–964.

56. Michaud M, Cognat V, Duchêne AM, Maréchal-Drouard L (2011) A global picture of tRNA genes in plant genomes. Plant J 66: 80–93.

57. Finn RD, Coggill P, Eberhardt RY, Eddy SR, Mistry J, Mitchell AL, Potter SC, Punta M, Qureshi M, Sangrador-Vegas A, et al. (2016) The Pfam protein families database: towards a more sustainable future. Nucleic Acids Res 44: D279–D285.

58. Rognes T, Flouri T, Nichols B, Quince C, Mahé F (2016) VSEARCH: a versatile open source tool for metagenomics. PeerJ 4: e2584.

59. Altschul SF, Gish W, Miller W, Myers EW, Lipman DJ (1990) Basic local alignment search tool. J Mol Biol 215: 403–410.

60. Galindo-González L, Mhiri C, Deyholos MK, Grandbastien MA (2017) LTR-retrotransposons in plants: Engines of evolution. Gene 626: 14–25.

61. Yang T, Poovaiah BW (2002) A calmodulin-binding/CGCG box DNA-binding protein family involved in multiple signaling pathways in plants. J Biol Chem 277: 45049–45058.

62. Abe H, Urao T, Ito T, Seki M, Shinozaki K, Yamaguchi-Shinozaki K (2003) Arabidopsis AtMYC2 (bHLH) and AtMYB2 (MYB) function as transcriptional activators in abscisic acid signaling. Plant Cell 15: 63–78.

63. Yang A, Dai X, Zhang W-H (2012) A R2R3-type MYB gene, OsMYB2, is involved in salt, cold, and dehydration tolerance in rice. J Exp Bot 63: 2541–2556.

64. Gómez-Porras JL, Riaño-Pachón D, Dreyer I, Mayer JE, Mueller-Roeber B (2007) Genome-wide analysis of ABA-responsive elements ABRE and CE3 reveals divergent patterns in Arabidopsis and rice. BMC Genomics 8: 260.

65. Timmerhaus G, Hanke ST, Buchta K, Rensing SA (2011) Prediction and validation of promoters involved in the abscisic acid response in physcomitrella patens. Mol Plant 4: 713–729.

66. Vishwakarma K, Upadhyay N, Kumar N, Yadav G, Singh J, Mishra RK, Kumar V, Verma R, Upadhyay RG, Pandey M, et al. (2017) Abscisic Acid Signaling and Abiotic Stress Tolerance in Plants: A Review on Current Knowledge and Future Prospects. Front Plant Sci 08: 1–12.

67. Sagi M, Fluhr R, Lips SH (1999) Aldehyde oxidase and xanthine dehydrogenase in a flacca tomato mutant with deficient abscisic acid and wilty phenotype. Plant Physiol 120: 571–578.

68. Parry AD, Neill SJ, Horgan R (1988) Xanthoxin levels and metabolism in the wild-type and wilty mutants of tomato. Planta 173: 397–404.

69. Burbidge A, Grieve TM, Jackson A, Thompson A, Mccarty DR, Taylor IB (1999) Characterization of the ABA-deficient tomato mutant notabilis and its relationship with maize Vp14. Plant J 17: 427–431.

70. Harrison E, Burbidge A, Okyere JP, Thompson AJ, Taylor IB (2011) Identification of the tomato ABA-deficient mutant sitiens as a member of the ABA-aldehyde oxidase gene family using genetic and genomic analysis. Plant Growth Regul 64: 301–309.

71. Maruyama KY, Todaka DA, Mizoi JU, Yoshida TA, Kidokoro SA, Matsukura SA, Takasaki HI, Sakurai TE, Yamamoto YOY, Yoshiwara KY (2012) Identification of Cis-Acting Promoter Elements in Cold- and Dehydration-Induced Transcriptional Pathways in Arabidopsis, Rice, and Soybean. DNA Res 19: 37–49.

72. Perlman PS, Boeke JD (2004) Ring around the retroelement. Science 303: 182–184.

73. Kilzer JM, Stracker T, Beitzel B, Meek K, Weitzman M, Bushman FD (2003) Roles of host cell factors in circularization of retroviral DNA. Virology 314: 460–467.

74. Li L, Olvera JM, Yoder KE, Mitchell RS, Butler SL, Lieber M, Martin SL, Bushman FD (2001) Role of the non-homologous DNA end joining pathway in the early steps of retroviral infection. EMBO J 20: 3272–3281.

75. Flavell AJ, Ish-Horowicz D (1981) Extrachromosomal circular copies of the eukaryotic transposable element copia in cultured Drosophila cells. Nature 292: 591–595.

76. Flavell AJ, Ish-horowicz D (1981) Extrachromosomal circular copies of the eukaryotic transposable element copia in cultured Drosophila cells. Nature 292: 591–595.

77. Tam SM, Causse M, Garchery C, Burck H, Mhiri C, Grandbastien M-A (2007) The distribution of copia-type retrotransposons and the evolutionary history of tomato and related wild species. J Evol Biol 20: 1056–1072.

78. Yin H, Liu J, Xu Y, Liu X, Zhang S, Ma J, Du J (2013) TARE1, a Mutated Copia-Like LTR Retrotransposon Followed by Recent Massive Amplification in Tomato. PLoS One 8: e68587.

79. Wang Y, Tang X, Cheng Z, Mueller L, Giovannoni J, Tanksley SD (2006) Euchromatin and pericentromeric heterochromatin: comparative composition in the tomato genome. Genetics 172: 2529–2540.

80. Cavrak V V., Lettner N, Jamge S, Kosarewicz A, Bayer LM, Mittelsten Scheid O (2014) How a Retrotransposon Exploits the Plant’s Heat Stress Response for Its Activation. PLoS Genet 10: e1004115.

81. Strand DJ, Mcdonald JF (1985) Copia is transcriptionally responsive to environmental stress. Nucleic Acids Res 13: 4401–4410.

82. Li Z, Peng R, Tian Y, Han H, Xu J, Yao Q (2016) Genome-Wide Identification and Analysis of the MYB Transcription Factor Superfamily in Solanum lycopersicum. Plant Cell Physiol 57: 1657–1677.

83. Seo PJ, Xiang F, Qiao M, Park J-Y, Lee YN, Kim S-G, Lee Y-H, Park WJ, Park C-M (2009) The MYB96 transcription factor mediates abscisic acid signaling during drought stress response in Arabidopsis. Plant Physiol 151: 275–289.

84. Zhu N, Cheng S, Liu X, Du H, Dai M, Zhou D-X, Yang W, Zhao Y (2015) The R2R3-type MYB gene OsMYB91 has a function in coordinating plant growth and salt stress tolerance in rice. Plant Sci 236: 146–156.

85. Sugimoto K, Takeda S, Hirochika H (2000) MYB-related transcription factor NtMYB2 induced by wounding and elicitors is a regulator of the tobacco retrotransposon Tto1 and defense-related genes. Plant Cell 12: 2511–2528.

86. Corem S, Doron-Faigenboim A, Jouffroy O, Maumus F, Arazi T, Bouché N (2018) Redistribution of CHH Methylation and Small Interfering RNAs across the Genome of Tomato ddm1 Mutants. Plant Cell 30: tpc.00167.2018.

87. Kravchik M, Damodharan S, Stav R, Arazi T (2014) Generation and characterization of a tomato DCL3-silencing mutant. Plant Sci 221–222: 81–89.

88. Panda K, Ji L, Neumann DA, Daron J, Schmitz RJ, Slotkin RK (2016) Full-length autonomous transposable elements are preferentially targeted by expression-dependent forms of RNA-directed DNA methylation. Genome Biol 17: 170.

89. McCue AD, Panda K, Nuthikattu S, Choudury SG, Thomas EN, Slotkin RK (2015) ARGONAUTE 6 bridges transposable element mRNA-derived siRNAs to the establishment of DNA methylation. EMBO J 34: 20–35.

90. Nuthikattu S, McCue AD, Panda K, Fultz D, DeFraia C, Thomas EN, Slotkin RK (2013) The Initiation of Epigenetic Silencing of Active Transposable Elements Is Triggered by RDR6 and 21-22 Nucleotide Small Interfering RNAs. Plant Physiol 162: 116–131.

91. Herr AJ, Jensen MB, Dalmay T, Baulcombe DC (2005) RNA polymerase IV directs silencing of endogenous DNA. Science (80-) 308: 118–120.

92. Kanno T, Huettel B, Mette MF, Aufsatz W, Jaligot E, Daxinger L, Kreil DP, Matzke M, Matzke AJM (2005) Atypical RNA polymerase subunits required for RNA-directed DNA methylation. Nat Genet 37: 761–765.

93. Cuerda-Gil D, Slotkin RK (2016) Non-canonical RNA-directed DNA methylation. Nat Plants 2: 16163.

94. Matzke M a, Mosher R a (2014) RNA-directed DNA methylation: an epigenetic pathway of increasing complexity. Nat Rev Genet 15: 394–408.

95. Deaton A, Bird A (2011) CpG islands and the regulation of transcription. Genes Dev 25: 1010–1022.

96. Ashikawa I (2001) Gene-associated CpG islands in plants as revealed by analyses of genomic sequences. Plant J 26: 617–625.

## SUPPLEMENTARY REFERENCES

Harkess, A. et al. The asparagus genome sheds light on the origin and evolution of a young y chromosome. Nat. Commun. 8, (2017).

Zou, C. et al. A high-quality genome assembly of quinoa provides insights into the molecular basis of salt bladder-based salinity tolerance and the exceptional nutritional value. Cell Res. 27, 1327–1340 (2017).

