## Supplementary figures and images for "Environmental and epigenetic regulation of *Rider* retrotransposons in tomato"

### Supplemental Figures

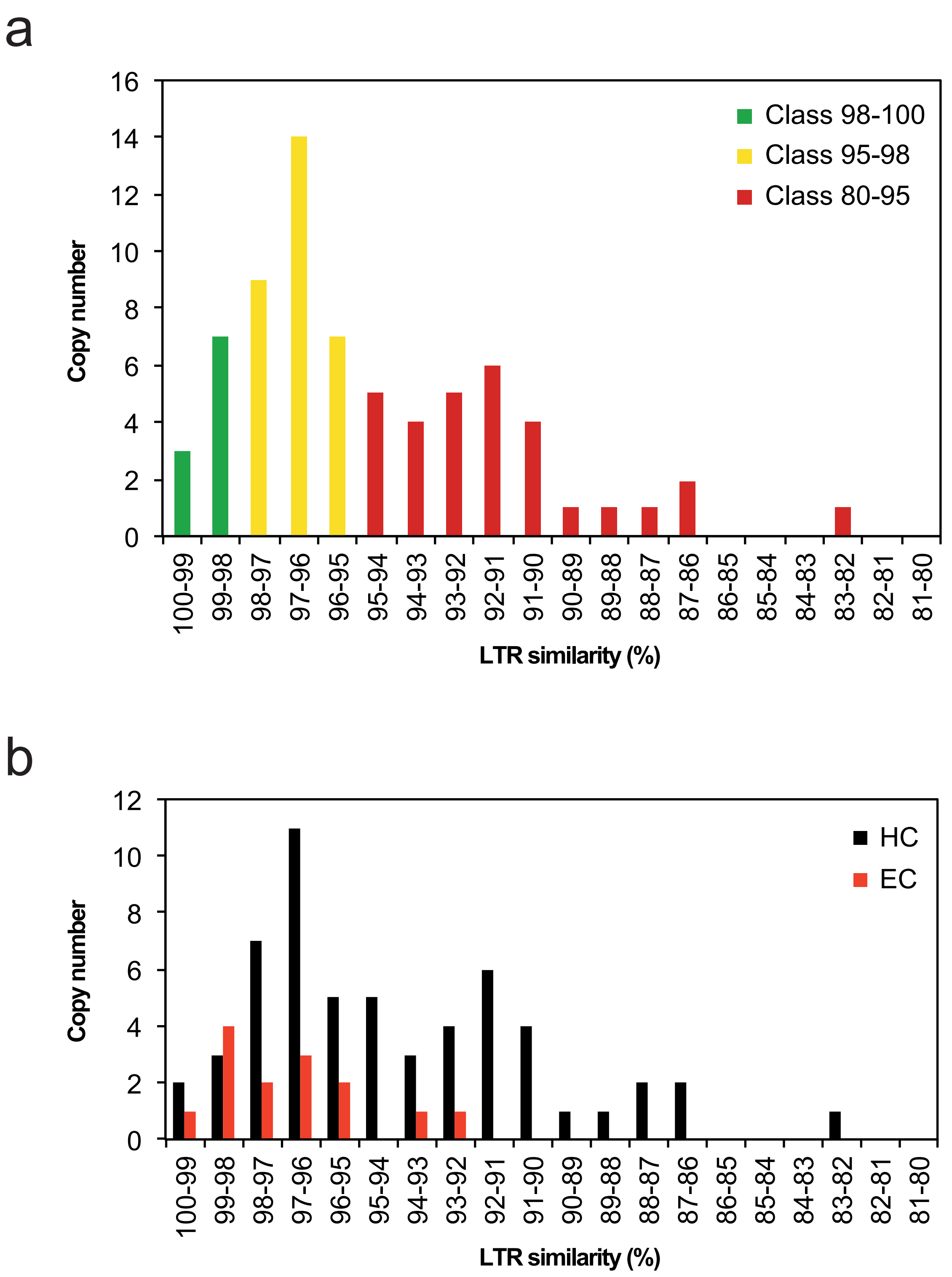

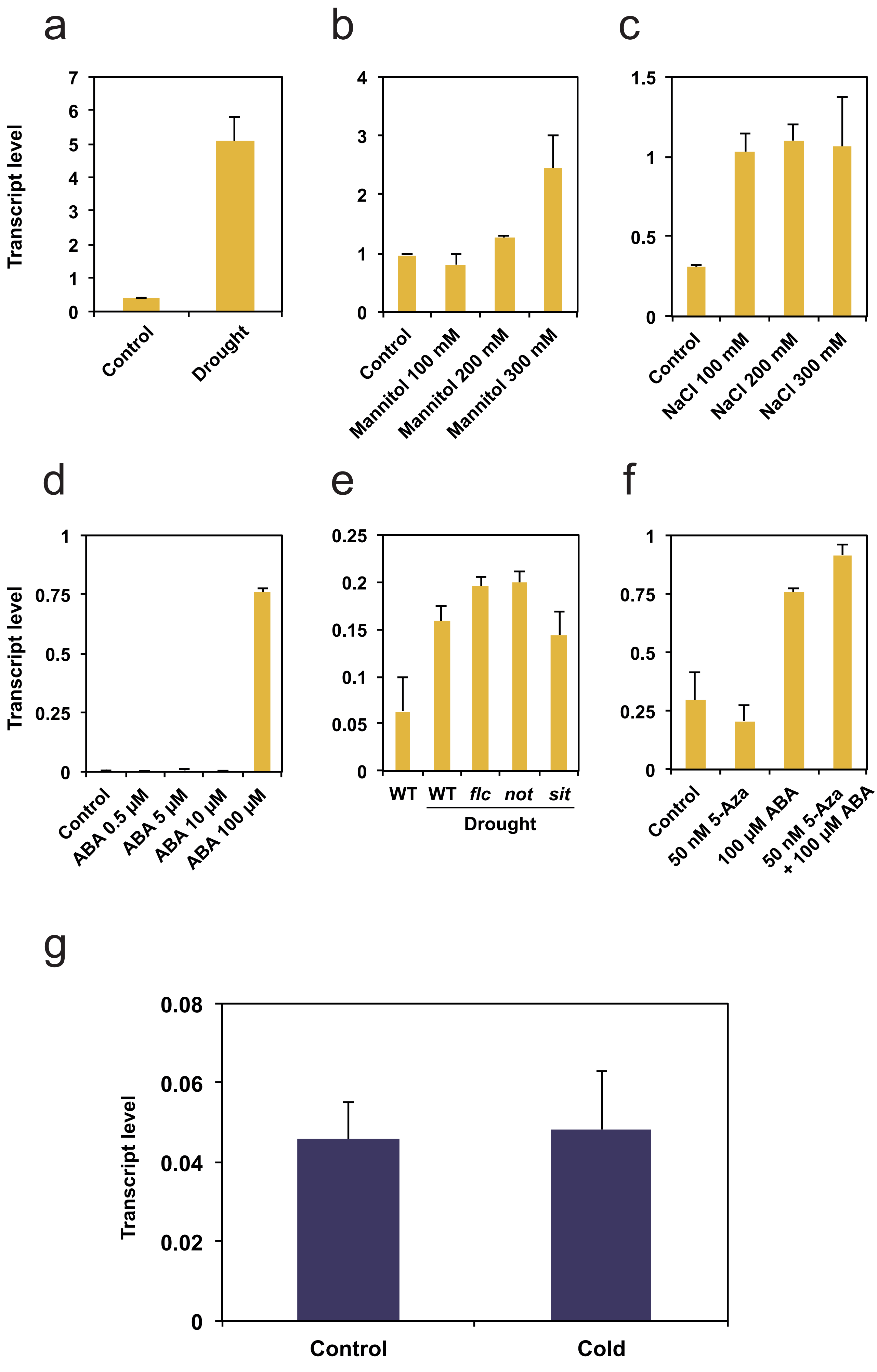

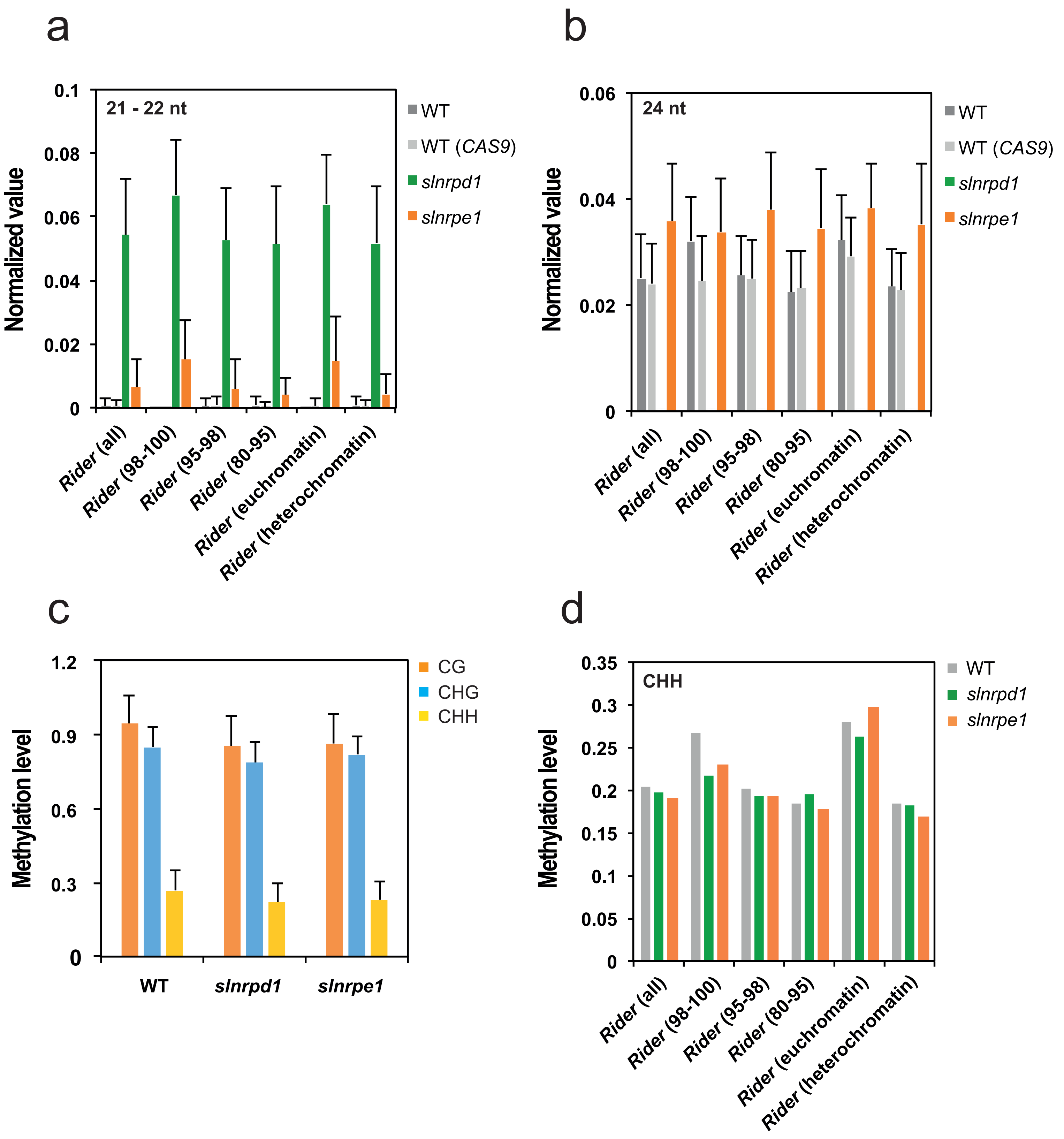

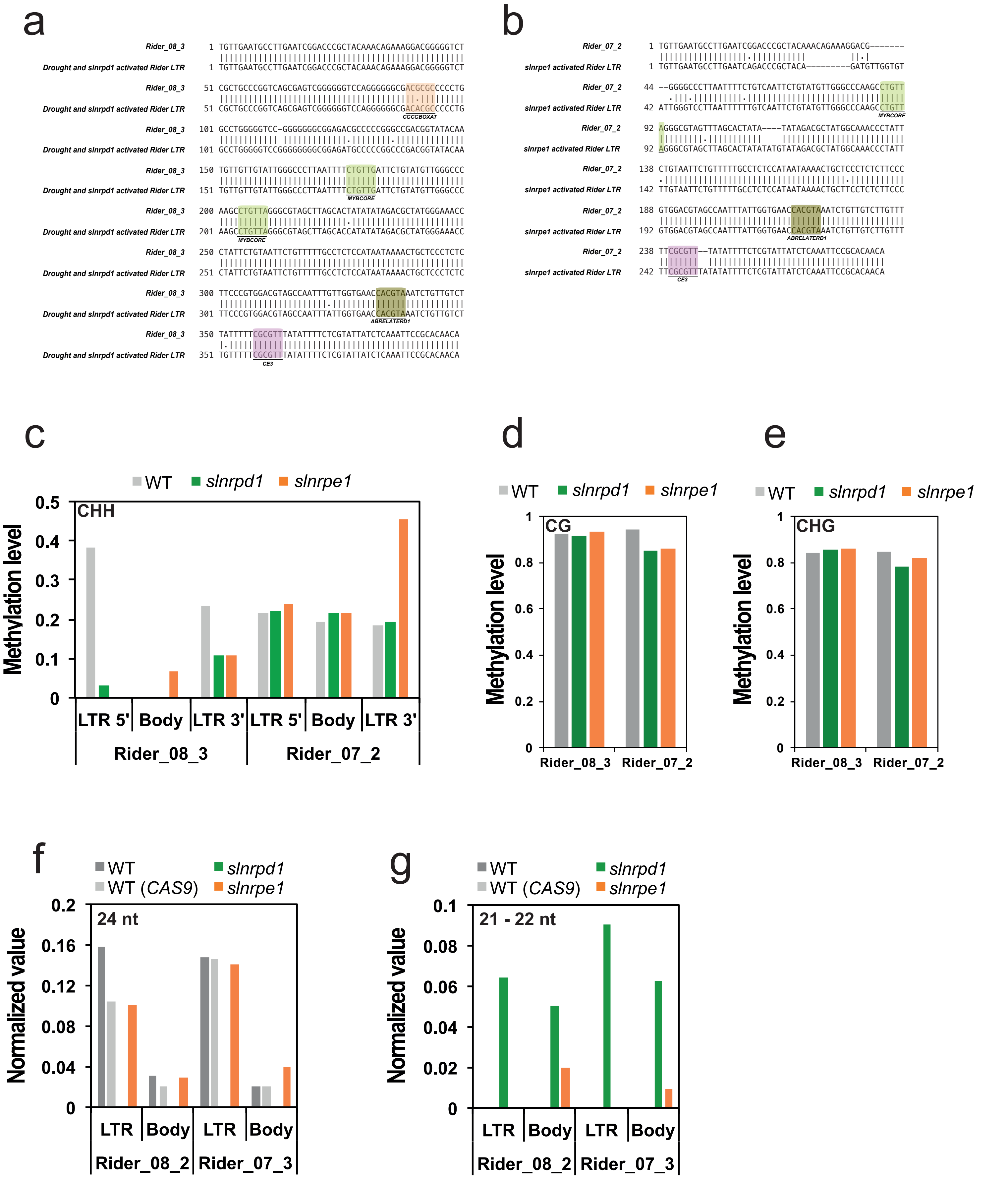

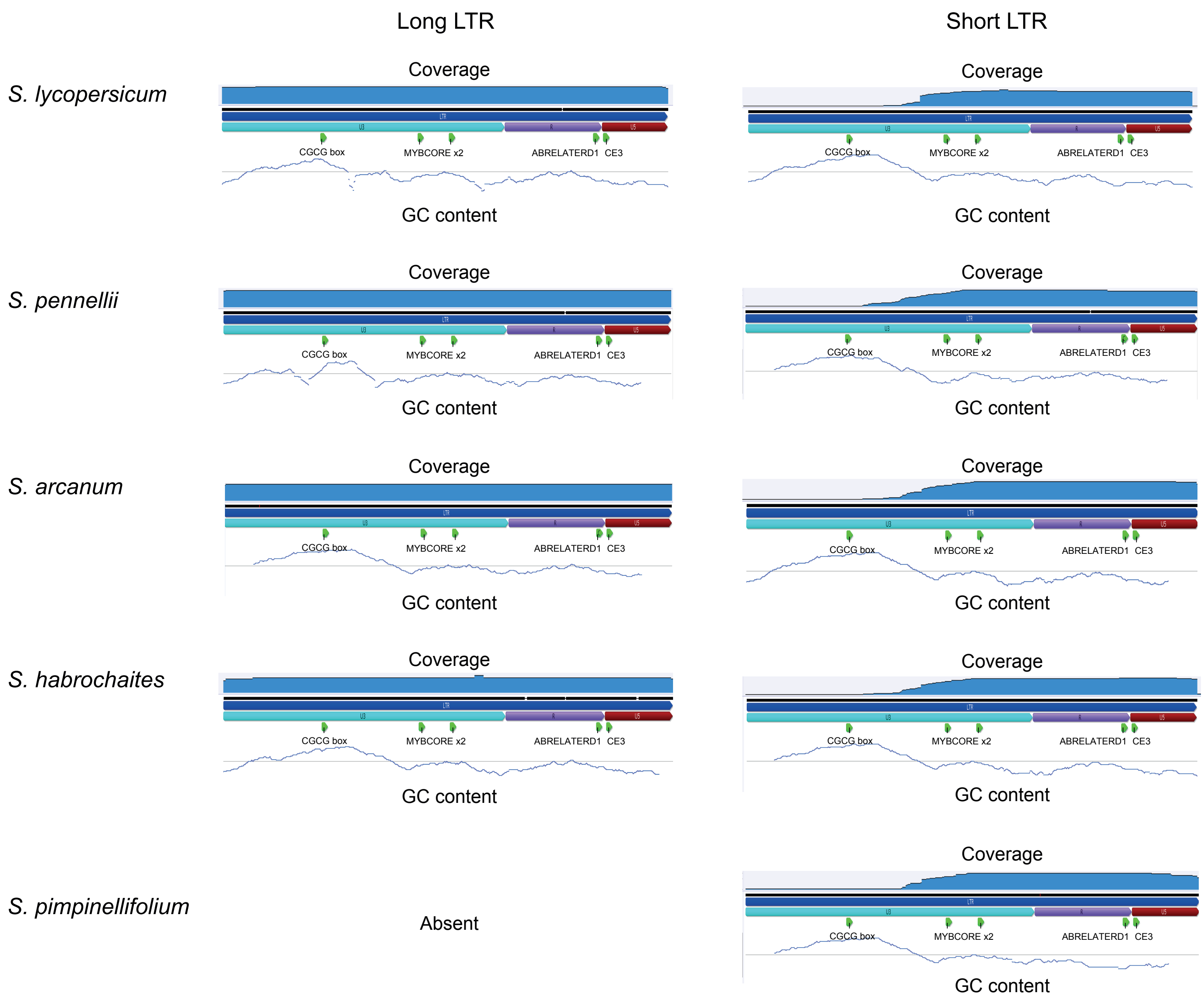

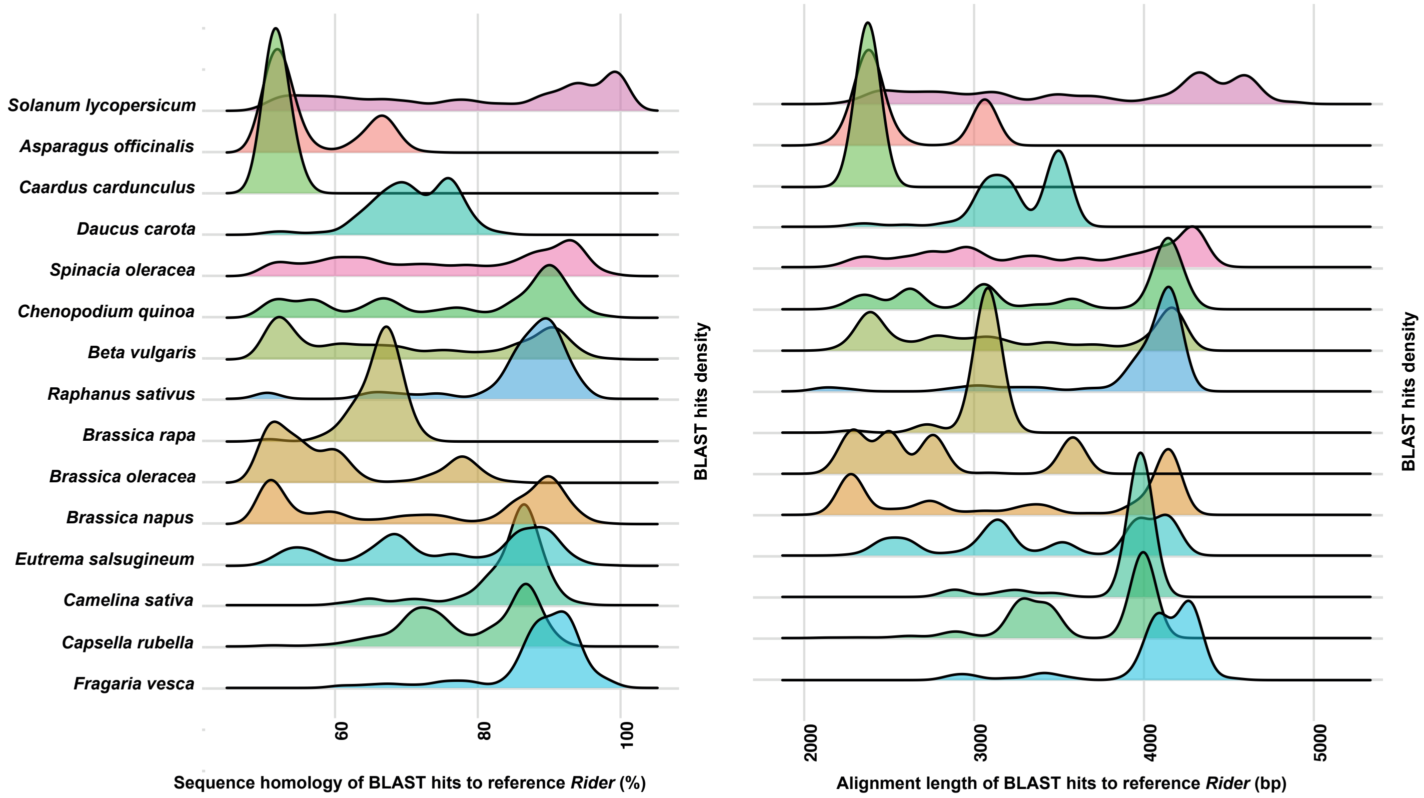

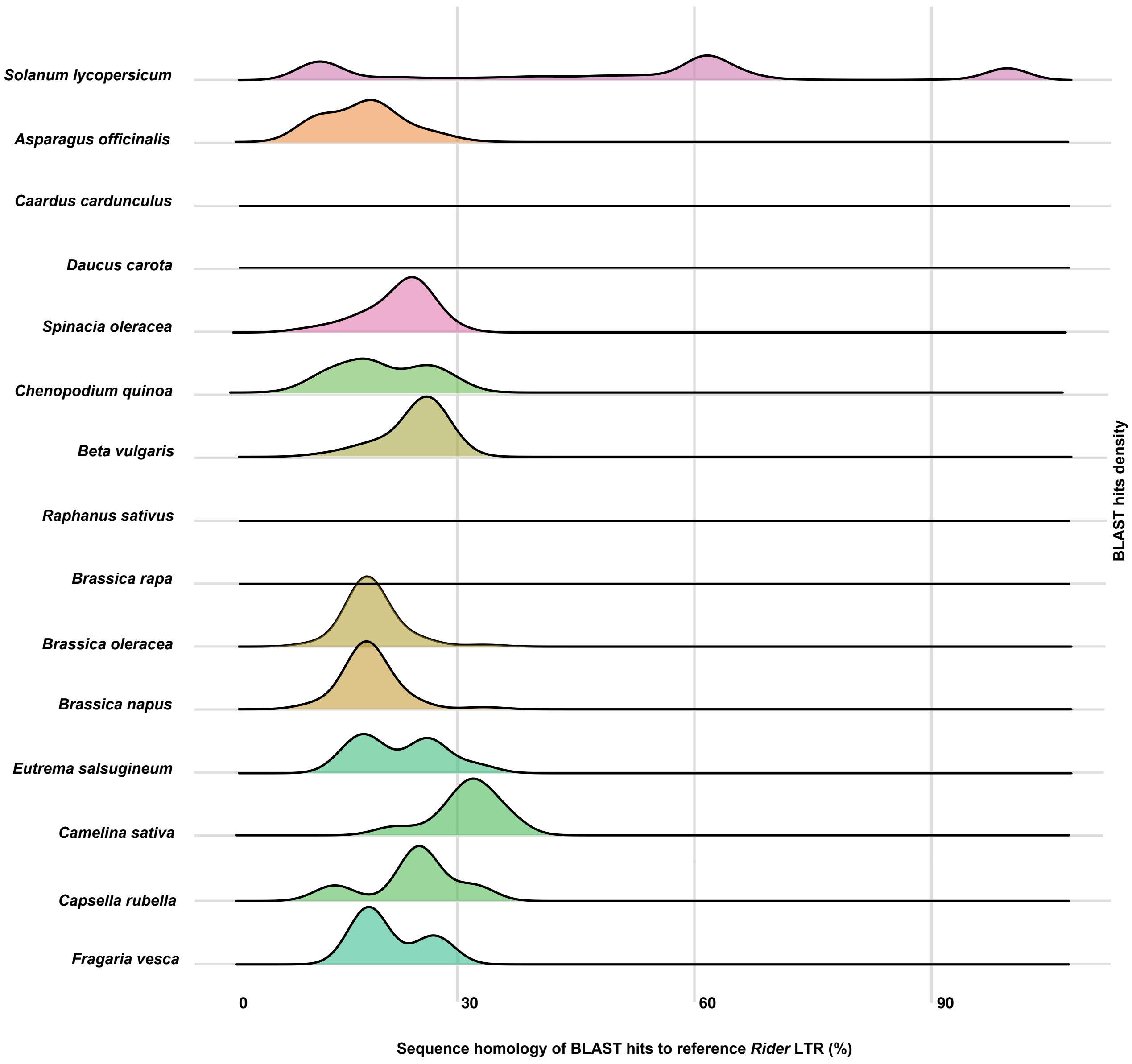
